# The OxyR-Responsive TacAT Module Promotes Oxidative Stress Adaptation and *In Vivo* Fitness in *K. pneumoniae*

**DOI:** 10.64898/2026.09.23.753711

**Authors:** Jun Zhou, Haichuan Ma, Zhou Sha, Shuzhi Liu, Jiayuan Rao, Ninglin Zhao, Xingyu Mou, Chunyi Chen, Hong Li, Huanxiang Liu, Haibo Wu, Rui Bao

**Author notes:** Corresponding authors: Rui Bao, E-mail addresses; Haibo Wu, E-mail addresses. The first three authors should be regarded as Joint First Authors.

## Abstract

During infection, *Klebsiella pneumoniae* must withstand host-derived oxidative stress not only by detoxifying reactive oxygen species (ROS) but also by repairing oxidative damage to cellular physiology. How these detoxification and stress-adaptive responses are coordinated remains unclear. Here, we identify a plasmid-encoded *tacAT* locus as an OxyR-responsive regulatory module that supports peroxide tolerance and in vivo fitness in *K. pneumoniae*. Guided by the conservation of the OxyR DNA-binding domain and its consensus binding motif, we surveyed promoter-proximal regions of the *K. pneumoniae CRK3022* chromosome and plasmids for candidate OxyR-responsive loci. This analysis identified *tacAT*, a putative type II toxin–antitoxin (TA) module, as the only plasmid-associated candidate. OxyR directly bound the *tacAT* promoter and contributed to *tacAT* induction during hydrogen peroxide (H_o_O_2_) stress. The TacAT complex exhibited DNA-binding and transcriptional regulatory activity, promoting the expression of genes linked to protein quality control and membrane/envelope homeostasis, including *clpB, htpG, cadC, bhsA*, and *marA*. Deletion of *tacAT* compromised survival under lethal peroxide challenge and attenuated bacterial dissemination, tissue pathology, and inflammatory responses in a murine bacteremia model. These findings define an OxyR–TacAT regulatory branch that connects peroxide sensing with stress-adaptive gene expression, revealing how a TA-associated module can be integrated into stress-adaptive programs during host-associated oxidative stress.

**Highlight:** Promoter motif screening identifies *tacAT* as a candidate OxyR-responsive TA-associated module OxyR directly binds the *tacAT* promoter and contributes to peroxide-induced *tacAT* expression TacAT promotes stress-response genes linked to protein quality control and envelope homeostasis TacAT supports oxidative stress survival and *in vivo* fitness of *Klebsiella pneumoniae*

**Graphical Abstract:** Proposed model of the OxyR-TacAT regulatory branch in *K. pneumoniae*. During infection, *K. pneumoniae* encounters host-derived oxidative stress. Oxidized OxyR activates classical antioxidant responses and contributes to *tacAT* induction by binding an OxyR-like motif in the *tacAT* promoter region. The induced TacAT complex binds OP1 within its own promoter and promotes the expression of stress-response genes, including *clpB* and *htpG*, which are associated with protein quality control, and *cadC*, *bhsA*, and *marA*, which are associated with membrane/envelope homeostasis. This regulatory branch supports bacterial survival under severe oxidative stress and contributes to in vivo fitness during systemic infection. Dashed arrows indicate proposed or indirect regulatory effects.

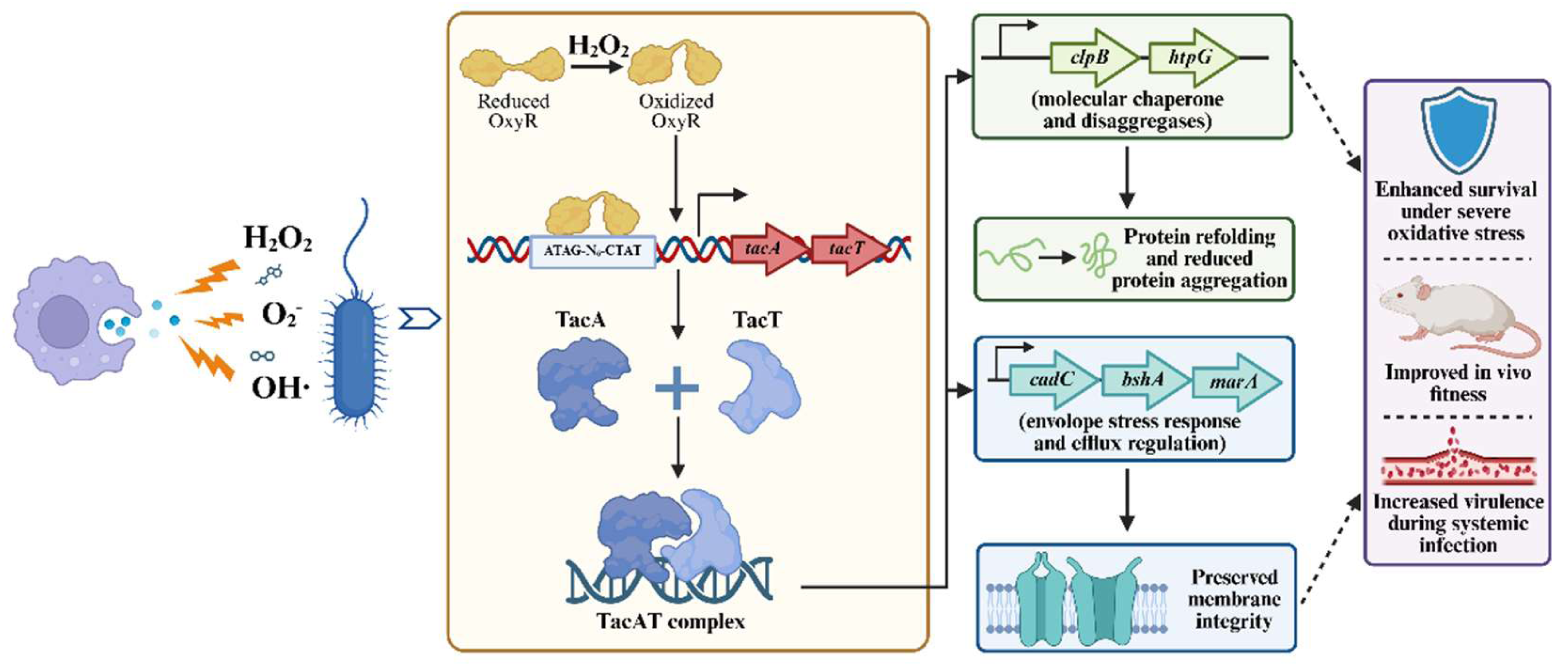

## Introduction

*Klebsiella pneumoniae* has emerged as a formidable public health threat, manifesting as a globally significant pathogen responsible for severe invasive infections, including bacteremia, pneumonia, and liver abscesses (1). While the proliferation of multidrug-resistant (2) and hypervirulent lineages (3) has precipitated a therapeutic impasse, clinical observations reveal that *K. pneumoniae* infections frequently exhibit recalcitrance and recurrence even following aggressive antimicrobial therapy (4). This phenomenon suggests that beyond genetic resistance, the pathogen possesses remarkable adaptive plasticity, allowing it to survive within the host environment. Accumulating evidence implicates the formation of stress-tolerant subpopulations as a pivotal driver of this chronic persistence (5). Unlike classical resistance, which relies on heritable mutations, this persistence strategy enables bacteria to withstand hostile conditions through transient physiological dormancy and metabolic remodeling (6), thereby sustaining infection reservoirs that are notoriously difficult to eradicate.

Upon invading the host, *K. pneumoniae* immediately encounters a hostile microenvironment characterized by intense oxidative stress. As a primary arm of innate immunity, host phagocytes generate a rapid oxidative burst, releasing lethal concentrations of ROS—such as superoxide anions, H_o_O_2_, and hydroxyl radicals—to eliminate invading pathogens (7). These highly reactive molecules indiscriminately damage bacterial DNA, proteins, and lipids (8), imposing a rigorous selection pressure on the pathogen. Consequently, the ability to rapidly sense and neutralize this oxidative assault is not merely a metabolic requirement but a fundamental determinant of pathogenicity and long-term survival. To cope with such extreme redox fluctuations, bacteria have evolved sophisticated defense networks that must coordinate immediate detoxification with deeper physiological adjustments to ensure viability under duress.

Central to the bacterial defense against ROS is the global transcriptional regulator OxyR, which senses H_o_O_2_ via redox-sensitive cysteine residues (9). Once activated, OxyR induces a classical antioxidant regulon that includes catalases, peroxidases, and other redox-balancing enzymes (10). These systems provide rapid detoxification of peroxide stress and are essential for maintaining redox homeostasis. However, during severe oxidative challenge, such as the oxidative burst encountered within phagocytes, detoxification alone may not be sufficient. ROS also damage proteins, membranes, and other cellular structures, creating a need for repair-oriented and stress-adaptive responses that act alongside enzymatic antioxidant defenses (11). This raises a broader regulatory question: beyond activating peroxide-detoxifying enzymes, does OxyR also connect redox sensing to downstream physiological adaptation?

Because OxyR homologs in Enterobacteriaceae share conserved DNA-binding features, motif-guided identification of OxyR-responsive promoters provides a useful strategy to explore potential direct outputs of the OxyR regulon. In this study, we first established the contribution of OxyR to severe peroxide stress survival in *K. pneumoniae CRK3022* and then surveyed promoter-proximal regions of the chromosome and plasmids for sequences resembling the conserved OxyR-binding motif. This search identified multiple candidate OxyR-responsive loci and unexpectedly highlighted a plasmid-encoded *tacAT* operon as the only plasmid-associated candidate. The *tacAT* locus is annotated as a putative type II TA module (12). TA systems are widely associated with bacterial stress tolerance, persistence, and growth modulation (13), yet their potential roles in stress-adaptive responses remain less well understood. Thus, *tacAT* represented a compelling candidate through which to examine whether OxyR-mediated peroxide sensing could be connected to TA-associated stress physiology and plasmid-encoded adaptive traits in *K. pneumoniae*.

Here, we show that OxyR directly binds the *tacAT* promoter and contributes to induced *tacAT* expression. We further demonstrate that the TacAT complex displays DNA-binding and transcriptional regulatory activity, promoting genes linked to protein quality control and membrane/envelope homeostasis. Rather than acting solely as a putative TA-associated growth-control module, TacAT supports survival under severe oxidative stress and contributes to *in vivo* fitness and virulence in a murine bacteremia model. Together, these findings reveal an OxyR–TacAT regulatory branch that connects redox sensing to stress-adaptive responses during infection.

## Results

### Motif-guided analysis identifies *tacAT* as a plasmid-associated candidate output of OxyR

OxyR has been implicated in H_2_O_2_ tolerance, biofilm formation, virulence, and antibiotic resistance in *K. pneumoniae* (14–16), but its contribution to survival under severe peroxide challenge has not been fully investigated. To examine this role in *K. pneumoniae CRK3022,* we constructed an *oxyR* deletion mutant (*ΔoxyR*) and quantified bacterial survival after exposure to 1% H_o_O_2_. The *ΔoxyR* strain showed a pronounced time-dependent loss of viability compared with the wild-type strain, with survival reduced by 72.58% after 20 min, 77.64% after 1 h, and 85.09% after 2 h of treatment (Fig. 1A). Thus, OxyR contributes substantially to survival under high-level peroxide stress in this strain.

**Figure 1.**
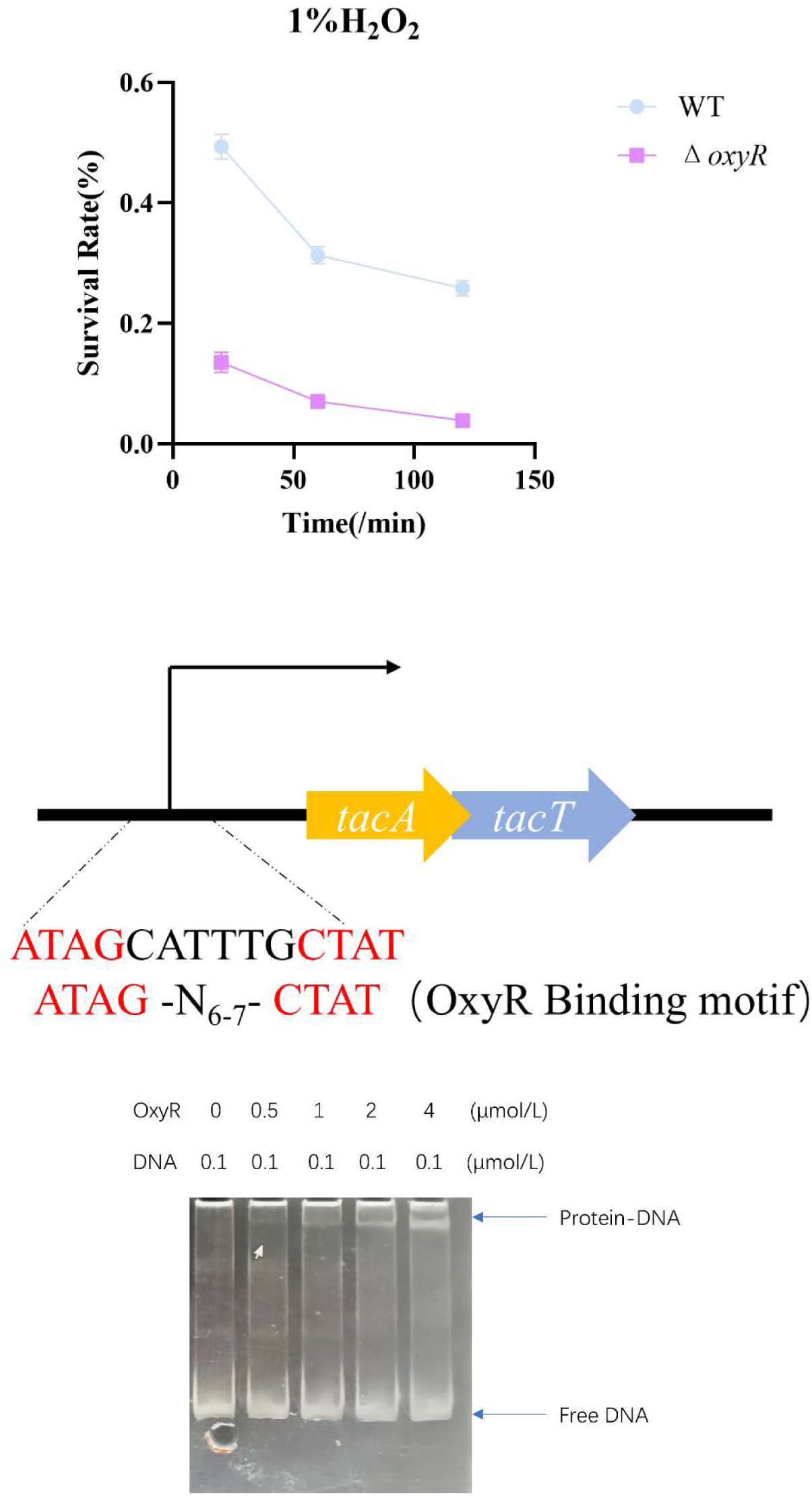
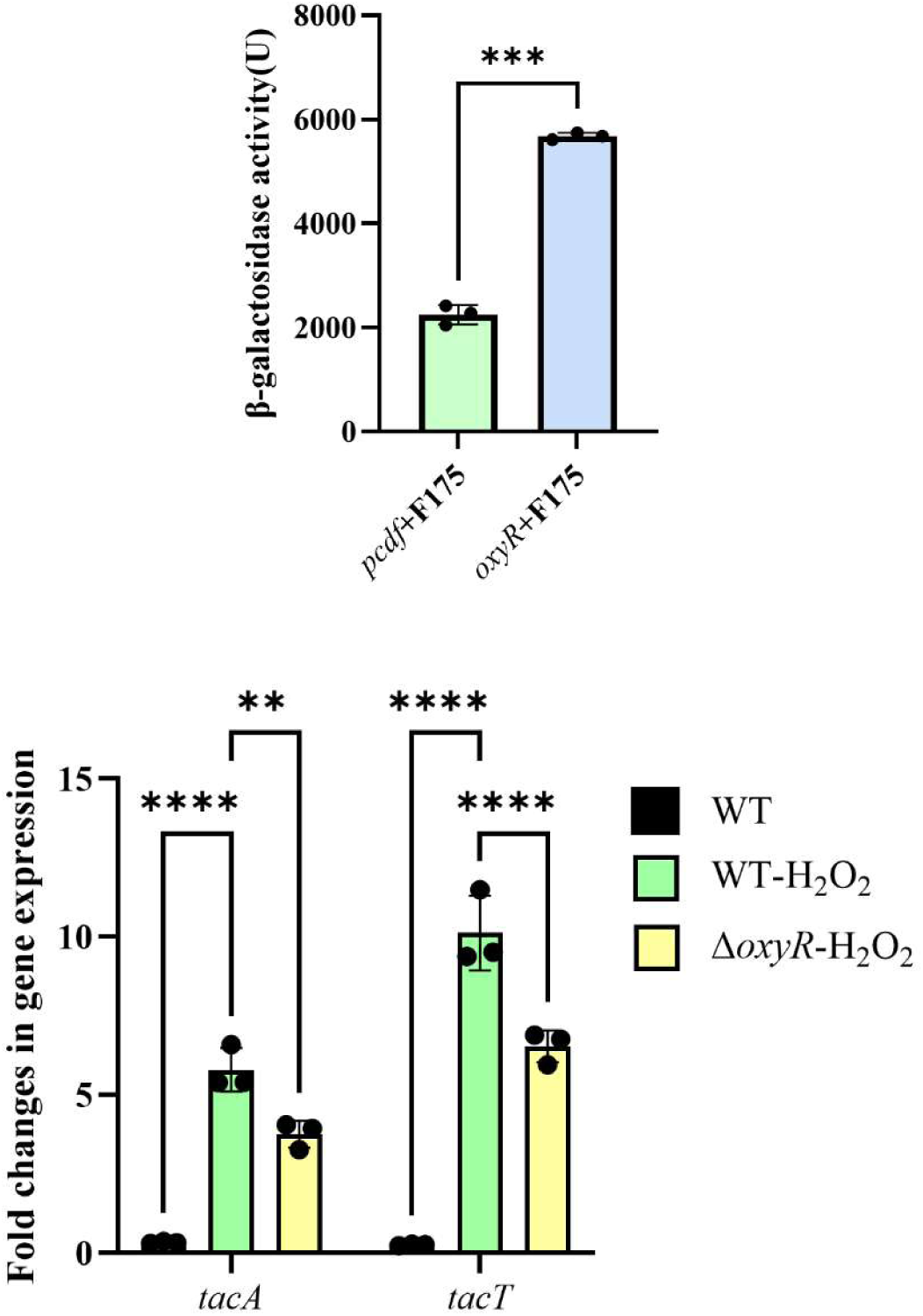
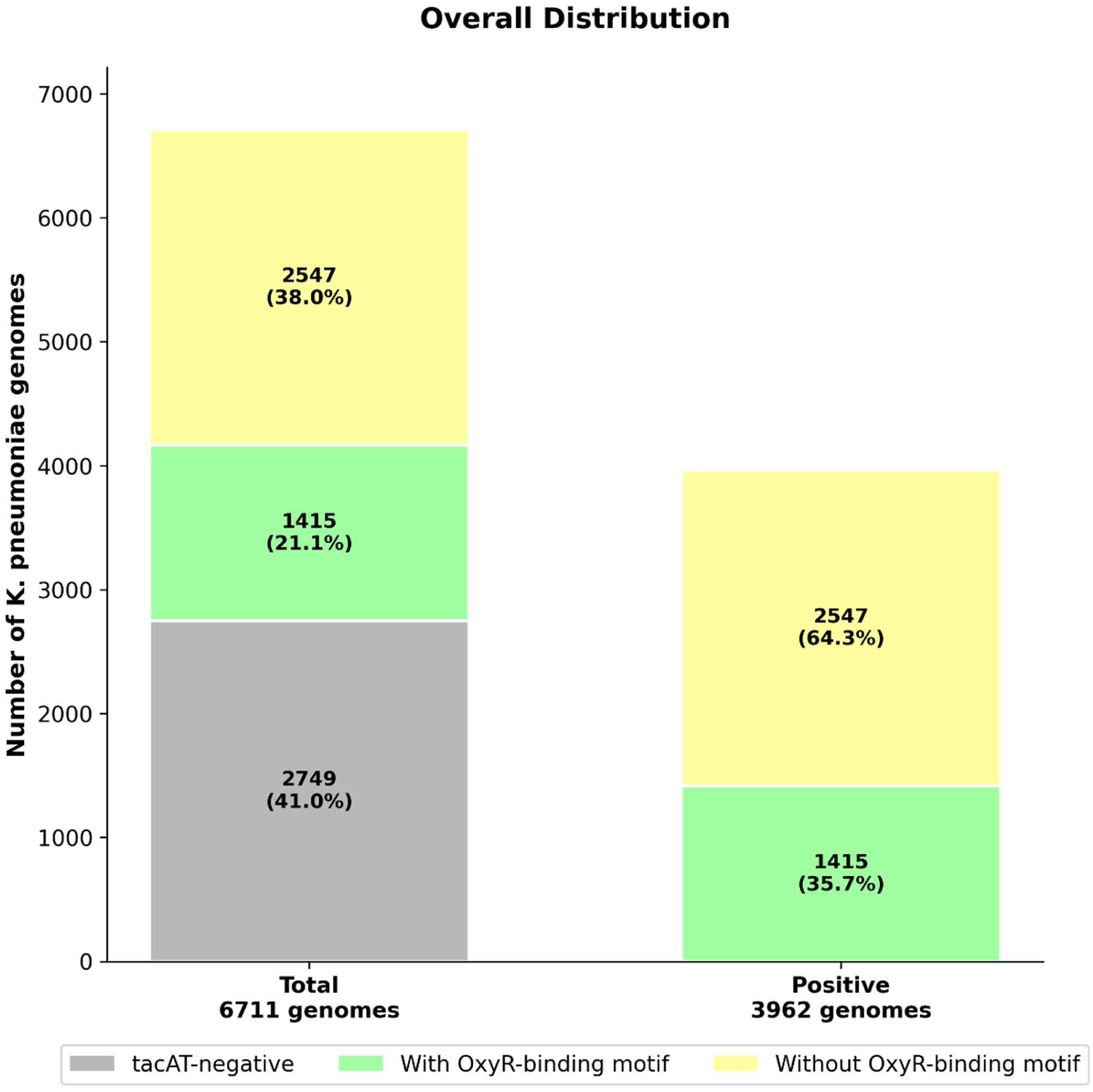
OxyR-dependent motif screening identifies *tacAT* as a candidate OxyR-responsive regulatory module. (A) Survival curves of *K. pneumoniae CRK3022* and Δ*oxyR* strains following exposure to 1% H_o_O_2_. (B) Alignment of the *E. coli* OxyR consensus motif with the OxyR-like sequence identified upstream of the *tacAT* operon. N denotes any nucleotide. (C) EMSA showing binding of OxyR to the *tacAT* noncoding region. (D) β-galactosidase reporter assay based on the intergenic region of the *tacAT* operon. F175 represents the native noncoding region of *tacAT*, pCDF represents the empty vector control, and OxyR represents the expression construct of *oxyR* cloned into the pCDF vector. (E) Relative transcription levels of *tacA* and *tacT* in wild-type and Δ*oxyR* strains under H_o_O_2_ stress, determined by RT-qPCR and normalized to *rpoD*. (F) Left stacked bar: distribution among all 6,711 publicly available *K. pneumoniae* genomes; 3,962 isolates (59.0%) carry intact *tacAT*, and 2,749 isolates (41.0%) are *tacAT*-negative. Right stacked bar: analysis restricted to the 3,962 *tacAT*-positive isolates, in which 1,415 genomes (35.7%) possess the putative OxyR-binding motif within the 100-bp upstream sequence of *tacAT*, and 2,547 genomes (64.3%) do not harbour this motif. Statistical significance was indicated as follows: **P < 0.01, *** P < 0.001, **** P < 0.0001, ns, not significant

We next asked which regulatory outputs might connect OxyR-mediated peroxide sensing to stress adaptation in *CRK3022*. OxyR DNA-binding specificity has been well characterized in *Escherichia coli*, where oxidized OxyR recognizes an inverted-repeat motif represented by ATAG-N6-7-CTAT (17). Comparison of *K. pneumoniae* OxyR with the *E. coli* ortholog showed strong conservation of the N-terminal DNA-binding region, including the helix–turn–helix domain (sequence identity, 100%; residues 18–38), and predicted structural models of the DNA-binding domain were highly similar (RMSD, 0.049 Å) (Fig. S1). These features supported the use of the *E. coli* OxyR motif as a guide to search for candidate OxyR-responsive loci in *K. pneumoniae*. We therefore performed a genome-wide survey of promoter-proximal noncoding regions in *CRK3022*, including both chromosomal and plasmid-borne sequences, to identify OxyR-like inverted repeats. This search identified 15 candidate OxyR-binding sites, including 14 chromosomal sites and one plasmid-associated site (Table S1 and Fig. S2). Several chromosomal candidates were linked to genes annotated in metal homeostasis, transport, stress regulation, and membrane-associated functions, categories consistent with known or plausible outputs of oxidative stress regulation. In contrast, the only plasmid-associated candidate mapped upstream of *tacAT*, a locus annotated as a putative type II TA module. This was unexpected because the candidate did not correspond to a canonical peroxide-detoxifying enzyme or redox-balancing protein. Instead, it suggested a possible plasmid-borne regulatory connection between OxyR and a TA-associated stress physiology module. The *tacAT*-associated motif was located in the intergenic region upstream of the operon and closely matched the OxyR consensus architecture (ATAG-CATTTG-CTAT) (Fig. 1B), making *tacAT* a compelling candidate for experimental validation.

To test whether OxyR directly recognizes the *tacAT* promoter, we first performed electrophoretic mobility shift (EMSA) assays using purified OxyR. OxyR shifted the *tacAT* promoter fragment in a concentration-dependent manner, indicating direct promoter binding (Fig. 1C). In parallel, β-galactosidase reporter assays showed that OxyR significantly increased *tacAT* promoter activity (Fig. 1D). OxyR-dependent promoter activation was detectable under the tested conditions (18).

We then examined *tacAT* expression during peroxide stress in *K. pneumoniae*. Under non-stress conditions, *tacAT* transcripts were detected at very low levels. Upon H2O_2_ exposure, *tacA* and *tacT* expression increased by approximately 17-fold and 38-fold, respectively, in the wild-type strain relative to the untreated control (Fig. 1E). In the Δ*oxyR* mutant, *tacAT* expression was reduced by approximately 35% compared with the wild type under the same condition. Thus, OxyR directly binds the *tacAT* promoter and contributes to maximal *tacAT* induction during peroxide stress. The residual induction observed in the Δ*oxyR* mutant further indicates that *tacAT* is not an exclusive output of OxyR but likely integrates additional regulatory inputs under oxidative stress.

To systematically assess whether the OxyR–TacAT regulatory linkage is broadly conserved across *K. pneumoniae*, we performed a large-scale genomic survey across a total of 6711 publicly available *K. pneumoniae* genomes. We first identified strains carrying an intact *tacAT* locus, yielding 3962 *tacAT*-positive isolates, accounting for 59.0% of the total analyzed strains (Fig. 1F). We then extracted the 100-bp noncoding sequence upstream of the *tacAT* translational start site from each positive strain and screened for the canonical OxyR-binding consensus motif (ATAG-N₆₋₇-CTAT).Across the 3962 *tacAT*-carrying strains, 1415 isolates (35.7%) harbored the OxyR-binding motif within the 100-bp upstream region (Fig. 1F). These findings indicate that an OxyR-compatible promoter architecture is present in a substantial subset of *tacAT*-positive *K. pneumoniae* genomes, suggesting that OxyR–TacAT regulatory coupling may extend beyond strain *CRK3022*.

### TacAT supports peroxide stress adaptation and exhibits OP1-dependent transcriptional regulatory activity

The *tacAT* locus identified in the OxyR motif screen is annotated as a putative type II TA-associated module, encoding a predicted antitoxin component, TacA, and a predicted toxin component, TacT. Because our study focuses on the stress-adaptive and regulatory functions of this module rather than the biochemical target of TacT toxicity, we first asked whether *tacAT* contributes to peroxide stress tolerance in *K. pneumoniae*. Deletion of the entire *tacAT* locus markedly impaired survival after exposure to 1% H_o_O_2_, with mutant showing a pronounced loss of viability compared with the wild-type strain after 2 h of treatment (Fig. 2A). Thus, an intact TacAT module is required for optimal survival under severe peroxide challenge.

**Figure 2.**
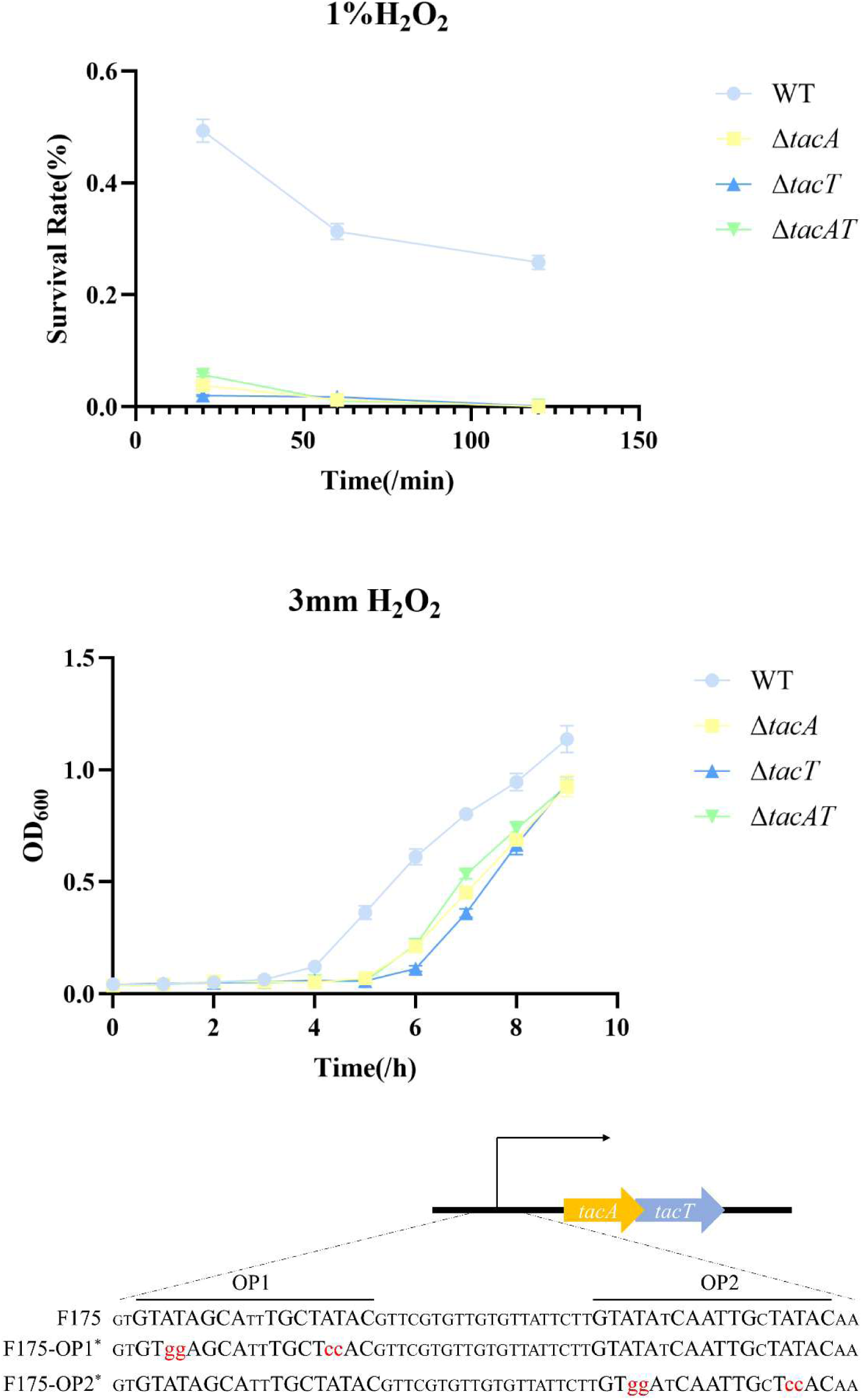
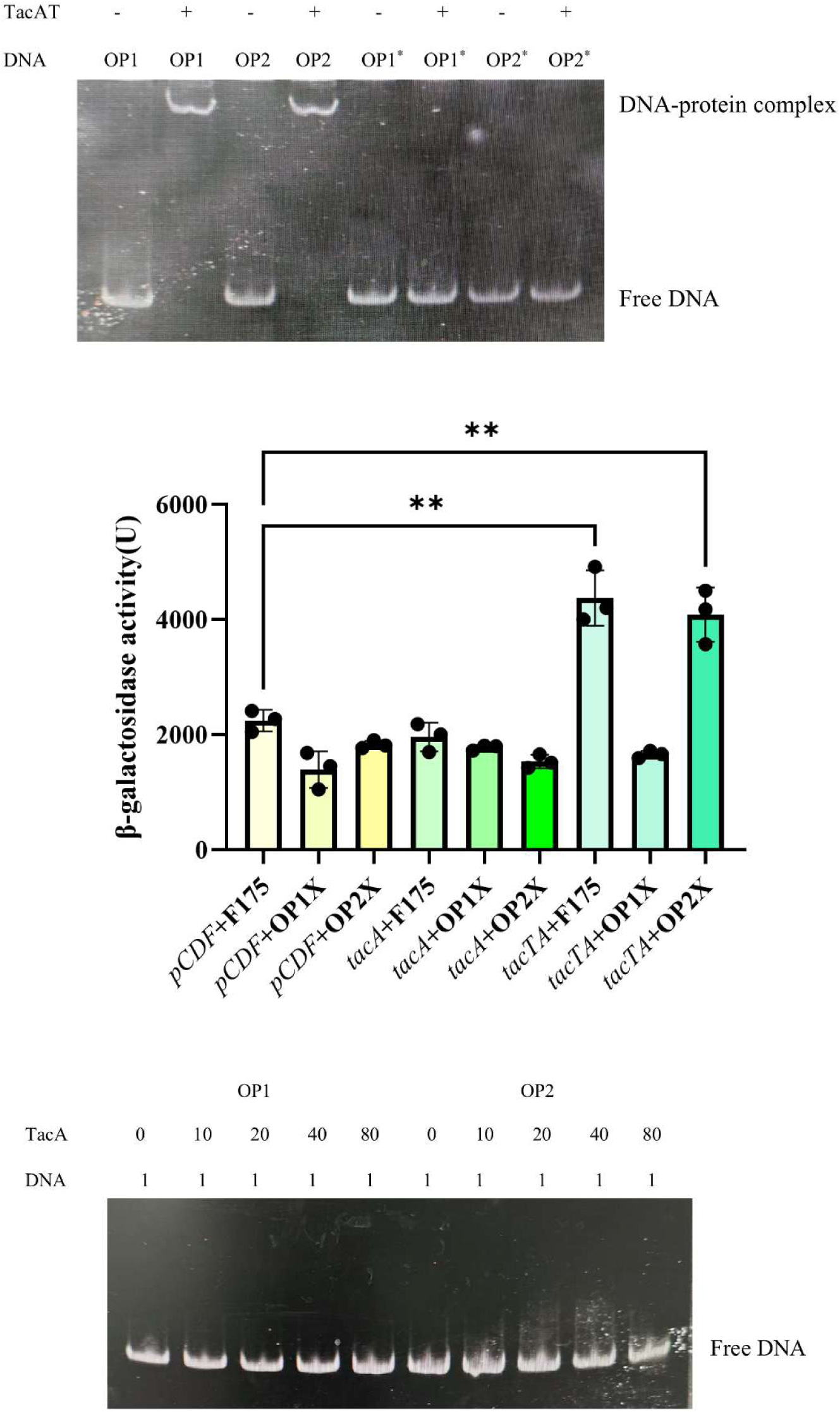
TacAT supports oxidative stress survival and promotes OP1-dependent *tacAT* promoter activity. (A) Survival curves of *K. pneumoniae CRK3022*, Δ*tacA,* Δ*tacT* and Δ*tacAT* strains following exposure to 1% H_o_O_2_. (B) Growth curves of *K. pneumoniae CRK3022*, Δ*tacA*, Δt*acT*, and Δ*tacAT* strains under sublethal oxidative stress (3 mM H_o_O_2_). (C) Nucleotide sequence of the *tacAT* intergenic region, showing two predicted pseudo-palindromic sequences (OP1 and OP2) and the corresponding nucleotide changes introduced in the mutant sequences (OP1* and OP2*). (D) EMSA using TacAT and the indicated DNA probes derived from OP1, OP2, and their corresponding mutated variants (OP1* and OP2*). (E) β-galactosidase reporter assays based on the *tacAT* intergenic region. F175 represents the native *tacAT* noncoding region, whereas OP1* and OP2* denote F175 derivatives carrying mutations in the first and second pseudo-palindromic sequences, respectively, as schematically illustrated in Fig. 2C. *pCDF* indicates the empty vector control, and *tacA* and *tacAT* correspond to expression constructs cloned into the *pCDF* vector. TacAT, but not TacA alone, enhanced promoter activity under the tested conditions. (F) EMSA using TacA and DNA probes corresponding to OP1 and OP2. Statistical significance was indicated as follows: **P < 0.01

TacAT also contributed to recovery from sublethal oxidative stress. In medium containing 3 mM H_o_O_2_, the wild-type strain resumed exponential growth after approximately 4 h, whereas the *tacAT* deletion strains showed a delayed recovery, entering exponential phase approximately 2 h later (Fig. 2B). This phenotype is consistent with impaired early adaptation and/or reduced viability during peroxide exposure. Together, the lethal-survival and sublethal-recovery assays indicate that TacAT is not merely associated with stress exposure at the transcriptional level but functionally contributes to peroxide stress adaptation.

We next examined whether this physiological role might be linked to promoter regulation by the TacAT complex. Many type II TA-associated antitoxins contain DNA-binding domains and regulate their cognate operons, most commonly through autorepression. Inspection of the *tacAT* intergenic region revealed two palindromic operator-like sequences, designated OP1 and OP2 (Fig. 2C) To test their contribution to TacAT-dependent regulation, the inverted repeats within OP1 or OP2 were individually mutated, generating OP1* (GTggAGCATTTGCTccAC) and OP2* (GTggATCAATTGCTccAC) promoter variants. EMSA analysis showed that mutation of OP1 disrupted TacAT binding to the corresponding probe, whereas the OP2 mutation had little effect on TacAT recognition under the tested conditions (Fig. 2D). Consistent with this binding pattern, β-galactosidase reporter assays showed that TacAT enhanced *tacAT* promoter activity, and this activation was abolished by mutation of OP1 but not OP2 (Fig. 2E). These results identify OP1 as the major operator required for TacAT-dependent promoter activation.

We then asked whether TacA alone was sufficient for this regulatory activity. Promoter activation was observed when TacA and TacT were co-expressed but not when TacA was expressed alone (Fig. 2E). In agreement with the reporter data, the TacAT complex showed detectable binding to operator probes, whereas TacA alone did not show detectable binding under the same EMSA conditions (Fig. 2D,F). These observations indicate that detectable TacA-associated DNA-binding activity requires TacT or assembly of the TacA–TacT complex under the tested conditions. Consistent with our previous single-knockout experiments, individual deletion of *tacA* or *tact* (Fig. 2A,B) severely impairs survival and recovery upon exposure to lethal concentrations of H_o_O_2_. The underlying mechanism remains to be defined, but a plausible working model is that TacT stabilizes TacA, promotes complex assembly, or favors a DNA-binding-competent conformation of TacAT. Thus, TacAT exhibits OP1-dependent positive regulatory activity of its own promoter under the tested conditions, differing from the autorepressive mode common to many type II TA systems.

### TacAT is associated with a stress-response transcriptional program linked to protein quality control and envelope adaptation

To identify stress-associated proteins whose abundance might be altered by TacAT, we performed comparative proteomic analysis as an exploratory candidate-generation approach. The Δ*tacAT* strain carrying pBAD-*tacAT* was compared with the Δ*tacAT* strain carrying the empty pBAD vector, allowing TacAT-dependent changes to be assessed in the same deletion background. This comparison revealed reproducible differences in global protein abundance between the two strains (Fig. 3A). Gene Ontology (GO) annotation of the differentially expressed proteins showed enrichment of the “response to stimulus” category within the biological process group (Fig. 3B), and multiple stress-associated proteins increased upon *tacAT* complementation (Table S2).Thus, restoration of TacAT expression was associated with broad remodeling of stress-related protein abundance, providing a candidate pool for identifying potential transcriptional outputs of the TacAT complex.

**Figure 3.**
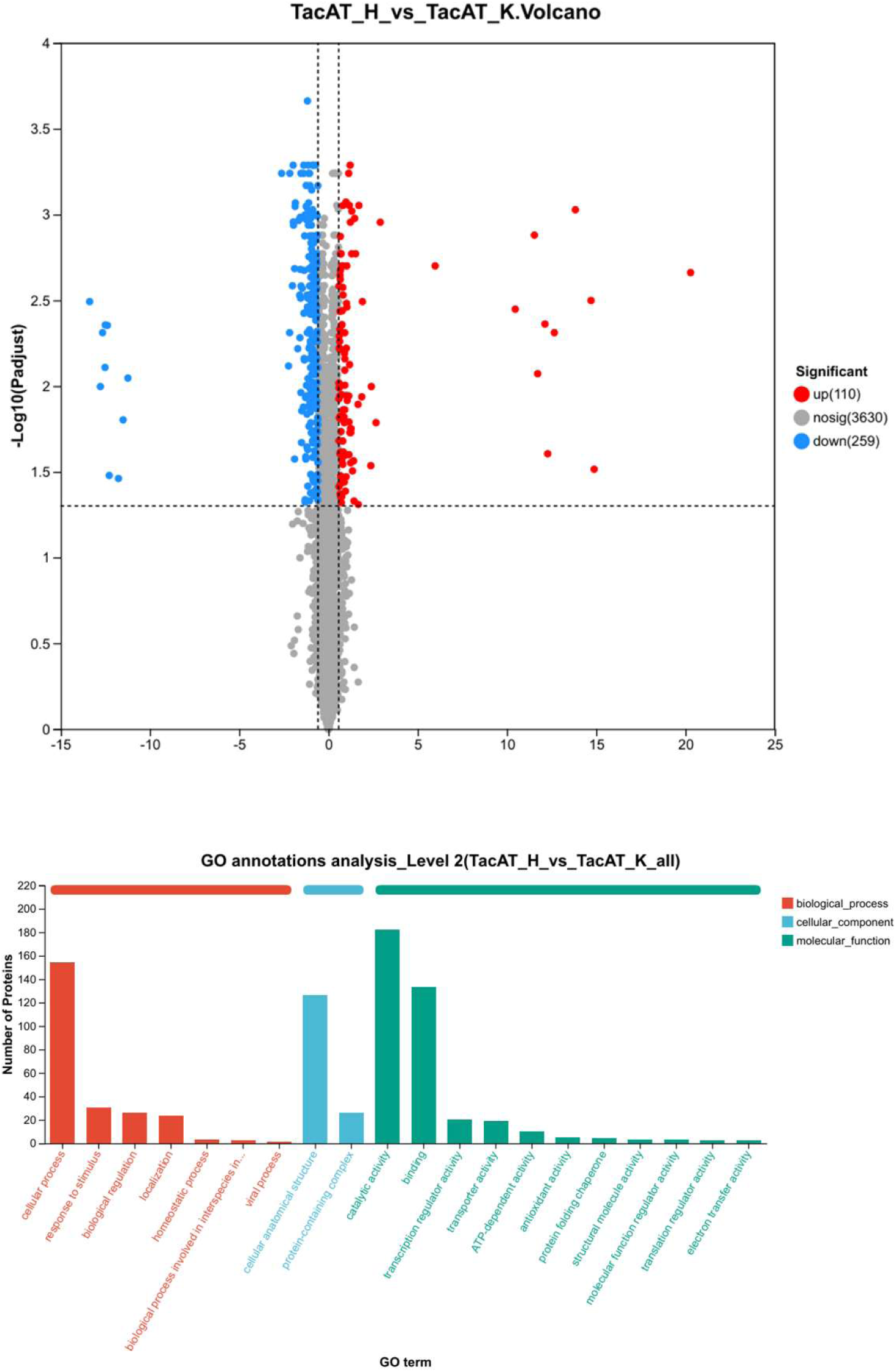
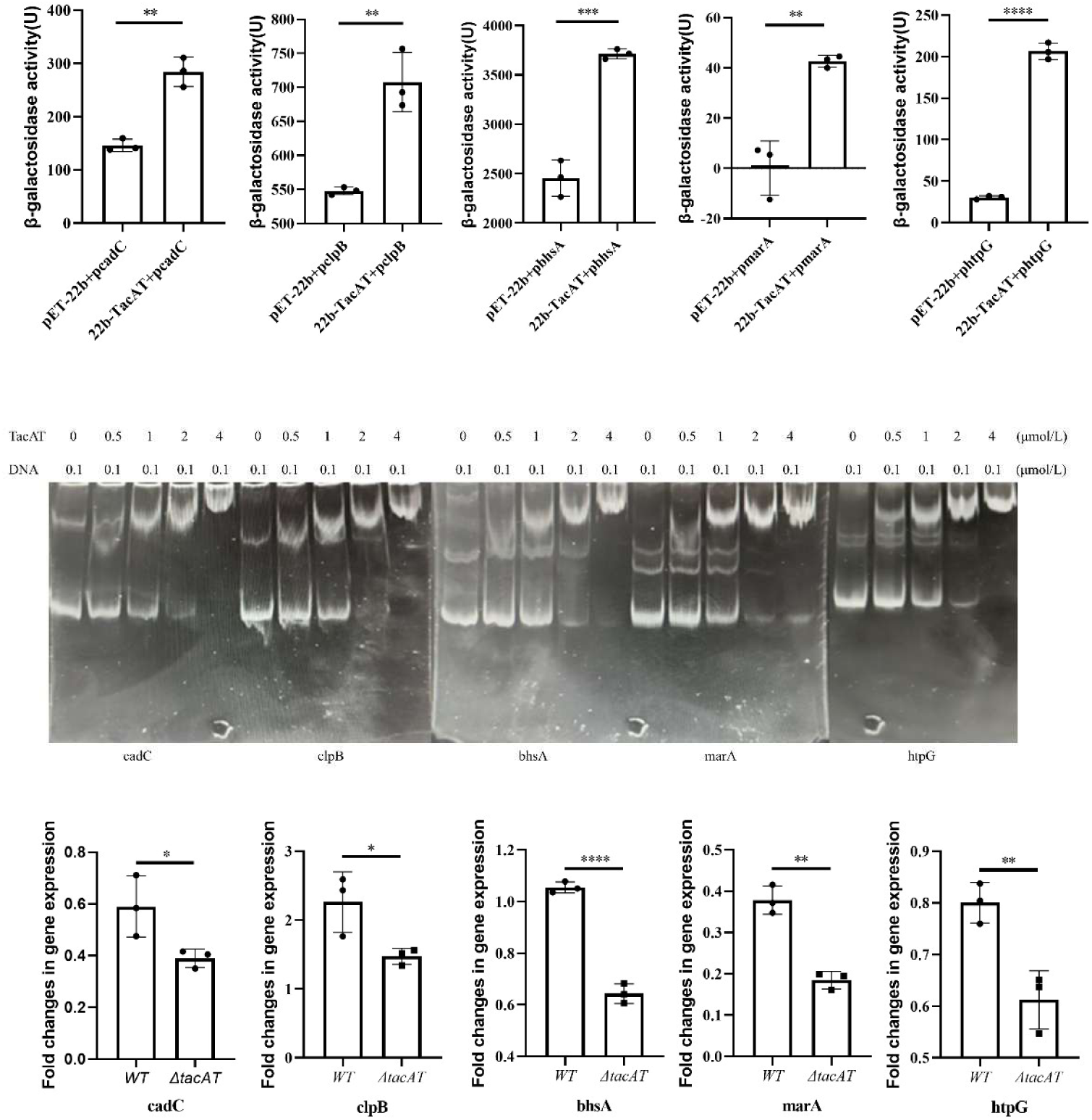
TacAT activates stress-response genes associated with protein quality control and membrane homeostasis. (A) Volcano plot showing differentially expressed proteins in Δ*tacAT*-pBAD-*tacAT* relative to Δ*tacAT*-pBAD. (B) GO enrichment analysis of differentially expressed proteins identified from the comparative proteomic analysis of Δ*tacAT*-pBAD-*tacAT* relative to Δ*tacAT*-pBAD. (C) β-galactosidase reporter assays measuring TacAT-dependent promoter activity of *cadC*, *clpB*, *bhsA*, *marA*, and *htpG*. (D) EMSA showing TacAT binding to the noncoding regions upstream of *cadC*, *clpB*, *bhsA*, *marA*, and *htpG*. (E) Transcriptional levels of *cadC*, *clpB*, *bhsA*, *marA*, and *htpG* in the wild-type and Δ*tacAT* strains under H_o_O_2_ stress. Statistical significance was indicated as follows: *P < 0.05, **P < 0.01, *** P < 0.001, **** P < 0.0001.

We next asked whether a subset of the TacAT-associated proteomic changes could be explained by direct promoter regulation. Stress-related genes that were upregulated upon *tacAT* complementation and represented distinct adaptive functions were selected for β-galactosidase reporter assays. This screen identified five promoters that responded significantly to TacAT: *clpB* and *htpG*, encoding chaperones involved in protein folding, refolding, and disaggregation (19, 20), and *cadC, bhsA,* and *marA,* which are linked to envelope stress adaptation, permeability control, or broader stress-responsive regulation (21–23) (Fig. 3C). Other tested stress-associated candidates did not show significant TacAT-dependent promoter activation under the same conditions (Fig. S3), indicating that TacAT responsiveness was restricted to a defined subset of loci rather than reflecting nonspecific activation of stress genes.

To determine whether TacAT can directly recognize the regulatory regions of these promoter-responsive genes, we performed EMSA using the corresponding upstream noncoding fragments. TacAT shifted the promoter-region probes of *clpB, htpG, cadC, bhsA*, and *marA*, supporting direct binding to these loci under the tested conditions (Fig. 3D). Together with the heterologous reporter assays, these data identify *clpB*, *htpG*, *cadC*, *bhsA*, and *marA* as candidate direct TacAT-responsive targets under the tested conditions. Functionally, these targets converge on two stress-adaptive themes: restoration of damaged or misfolded proteins through chaperone-associated protein quality control, and protection of the cell envelope through regulators or factors linked to membrane/envelope homeostasis.

We further examined whether TacAT is required for induction of these genes during peroxide stress in *K. pneumoniae.* Under H_o_O_2_ challenge, the mRNA levels of *clpB, htpG, cadC, bhsA*, and *marA* were significantly lower in the Δ*tacAT* mutant than in the wild-type strain (Fig. 3E). Thus, the TacAT-responsive targets identified through proteomics, reporter assays, and EMSA are also TacAT-dependent during oxidative stress. Together, these findings indicate that TacAT promotes a focused transcriptional program that links peroxide stress adaptation to protein quality control and envelope-associated protective functions.

### TacAT contributes to in vivo fitness during systemic infection in a murine bacteremia model

Having shown that TacAT supports peroxide stress adaptation and activates stress-response genes *in vitro*, we next asked whether this module contributes to bacterial fitness during systemic infection. Mice were infected intravenously with the wild-type strain or Δ*tacAT* strains using complementary high-dose and low-dose bacteremia models. In the high-dose challenge model, infection with 5 × 10⁶ CFU of the wild-type strain caused rapid mortality, with all mice succumbing by day 3 (Fig. 4A). By contrast, Δ*tacAT* infection significantly attenuated lethality, and 20–40% of infected mice survived until day 5. These results indicate that loss of *tacAT* reduces lethality of *K. pneumoniae* in systemic infection.

**Figure 4.**
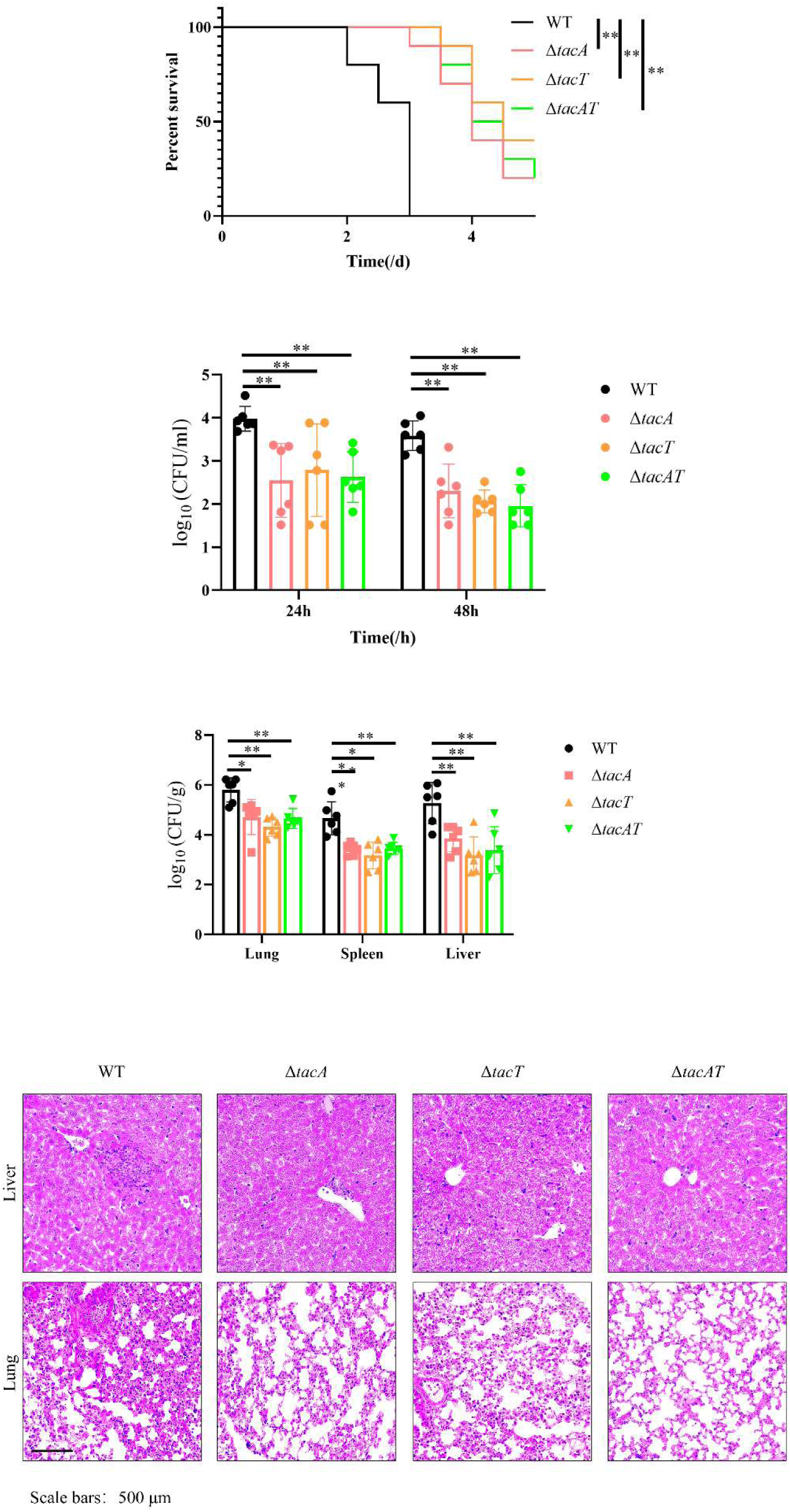
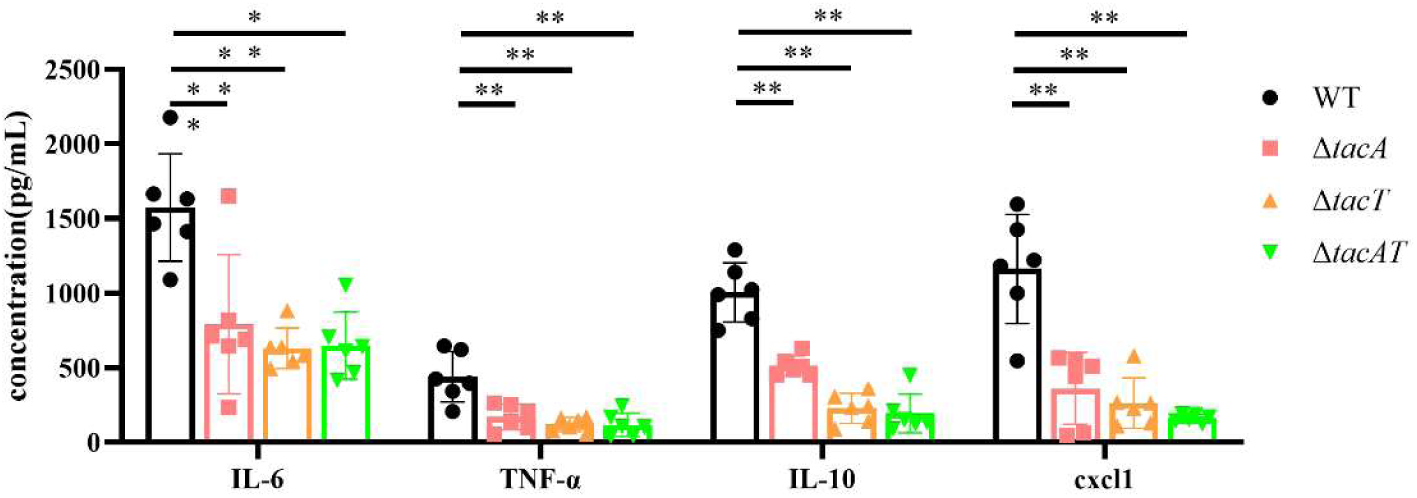
TacAT contributes to in vivo fitness during systemic infection in *K. pneumoniae*. (A) Survival curves of mice infected with a high bacterial dose. (n = 10 mice) (B) Bacterial loads in the blood of mice infected with a low bacterial dose. (n = 6 mice) (C) Bacterial burdens in major organs (lungs, spleen, and liver) at 48 h post-infection following low-dose challenge. For bacterial burden analyses, CFU values were log₁₀-transformed before visualization and statistical analysis. (n = 6 mice) (D) H&E staining of lung and liver tissues from mice infected with a low bacterial dose. (E) Serum levels of inflammatory cytokines (IL-6, TNF-α, IL-10, and CXCL1) at 48 h post-infection following low-dose challenge. (n = 6 mice) Statistical significance was indicated as follows: *P < 0.05, **P < 0.01.

We next used a low-dose infection model to determine whether this attenuation was associated with impaired bacterial persistence and dissemination. Following infection with 1 × 10⁵ CFU, Δ*tacAT* strains showed markedly reduced bacterial loads in blood, with 75–92% lower burdens at 24 h and 90–96% lower burdens at 48 h compared with the wild-type strain (Fig. 4B). At 48 h, bacterial burdens in the lungs, spleen, and liver were also reduced by 90–97%, 97–98%, and 96– 98%, respectively, in the Δ*tacAT* groups (Fig. 4C). Thus, *tacAT* is required for optimal survival and systemic spread *in vivo*.

The reduction in bacterial burden was accompanied by decreased tissue pathology. Wild-type infection caused pronounced inflammatory damage in the liver and lungs, including hepatic inflammatory infiltration and extensive pulmonary consolidation, septal thickening, and neutrophil-rich inflammatory infiltrates. In contrast, tissues from Δ*tacAT*-infected mice showed markedly attenuated pathology, with largely preserved lung architecture, open alveolar spaces, and limited inflammatory infiltration(Fig. 4D). Consistent with the reduced bacterial burdens and tissue damage, reduced bacterial burden was accompanied by lower circulating cytokine concentrations. Serum levels of IL-6, IL-10, TNF-α, and CXCL1 were significantly reduced compared with wild-type infection, with decreases of 49–59%, 61–73%, 48–80%, and 68–86%, respectively (Fig. 4E). These host-response readouts further support the attenuated *in vivo* fitness of Δ*tacAT* strains.

Together, these findings extend the *in vitro* function of TacAT to the infection setting: a module that promotes peroxide stress adaptation and stress-gene expression also supports bacterial persistence, tissue colonization, and inflammatory pathology in vivo. Although the specific host stresses responsible for this attenuation remain to be defined, the data are consistent with TacAT acting as a stress-adaptive fitness module during systemic infection.

## Discussion

This study identifies TacAT as an OxyR-responsive regulatory branch that links peroxide sensing to stress-adaptive gene expression in *K. pneumoniae*. Rather than representing an isolated TA-associated locus, our findings position *tacAT* within a broader redox-responsive regulatory framework. Motif-guided analysis of the *CRK3022* chromosome and resident plasmid replicons identified multiple candidate OxyR-responsive loci, among which *tacAT* was the only plasmid-associated candidate. OxyR directly bound the *tacAT* promoter and contributed to maximal peroxide-induced *tacAT* expression, whereas deletion of *tacAT* impaired survival under severe oxidative stress and attenuated systemic infection in mice. Together, these findings support a model in which TacAT may function as a stress-adaptive component of the OxyR response, complementing canonical peroxide detoxification pathways rather than replacing them.

OxyR has long been recognized as a central bacterial peroxide sensor that activates catalases, peroxidases, thiol-redox systems, and other antioxidant defenses (11). This detoxification-centered model explains how bacteria rapidly reduce ROS burden and restore redox balance. However, severe oxidative stress also causes secondary cellular damage, including protein oxidation, misfolding, membrane injury, and disruption of envelope integrity (24). Our findings suggest that in *K. pneumoniae*, OxyR-dependent adaptation extends beyond classical antioxidant enzymes to include regulatory modules involved in downstream damage management. The partial reduction of *tacAT* induction in the Δ*oxyR* mutant is particularly informative, indicating that OxyR acts as a major contributor but not the sole regulator of *tacAT* expression during peroxide stress. Such partial dependency is consistent with the architecture of bacterial stress networks, in which multiple regulatory pathways integrate redox state, envelope stress, metabolism, and growth conditions to fine-tune adaptive responses (25). Therefore, the OxyR–TacAT relationship should not be viewed as a simple linear pathway, but rather as one regulatory branch embedded within a multilayered oxidative stress response network.

To determine whether this regulatory relationship extends beyond the *CRK3022* strain, we surveyed 6,711 publicly available *K. pneumoniae* genomes. An intact *tacAT* locus was identified in 59.0% of isolates (3,962/6,711), and among these *tacAT*-positive strains, 35.7% retained a canonical OxyR-binding motif (ATAG-N₆₋₇-CTAT) within the 100-bp upstream noncoding region. This partial conservation pattern indicates that an OxyR-compatible promoter architecture is present in a subset of *tacAT*-positive genomes, suggesting that OxyR–*tacAT* regulatory coupling may represent a lineage-variable adaptive feature. The incomplete distribution of the motif is consistent with the dynamic nature of bacterial regulatory networks, in which newly acquired or pre-existing genetic modules may be differentially integrated into host-associated stress responses depending on genomic background and ecological selective pressure.

The presence of an OxyR-responsive motif in a substantial proportion of *tacAT*-positive strains supports the possibility that TacAT-mediated stress adaptation may provide a selective advantage under oxidative stress conditions. Canonical OxyR targets, including catalases and peroxidases, function as a first layer of defense by directly eliminating ROS and restoring redox homeostasis (11). In contrast, TacAT appears to represent a downstream adaptive layer that mitigates the secondary cellular damage caused by oxidative injury. Through activation of chaperone-associated factors such as *clpB* and *htpG* (19, 20), TacAT may promote recovery from protein oxidation and aggregation, while regulation of *cadC*, *bhsA*, and *marA* may contribute to envelope stabilization and stress tolerance (21–23). Thus, the OxyR–TacAT axis may complement classical antioxidant defenses by coordinating ROS detoxification with subsequent stress-adaptive responses.

The uneven distribution of the OxyR motif among *tacAT*-positive genomes further highlights the evolutionary flexibility of mobile genetic elements. Although plasmids are traditionally viewed as vehicles for antibiotic resistance and virulence dissemination, they can also serve as reservoirs of adaptive regulatory modules that enhance bacterial survival in fluctuating environments (26, 27). In strain *CRK3022*, the plasmid localization of *tacAT* exemplifies how horizontally acquired TA modules can be functionally rewired to plug into conserved host stress regulatory networks. Variation within promoter regions of horizontally acquired genes provides ongoing opportunities for regulatory rewiring, allowing accessory modules to acquire new connections with core stress pathways. Further comparative genomic and evolutionary analyses will be required to determine how frequently such regulatory integration occurs across *K. pneumoniae* populations.

A notable feature of *K. pneumoniae* TacAT is its OP1-dependent positive regulatory activity. Many type II TA systems autoregulate their own operons through antitoxin-mediated repression, often influenced by TA stoichiometry and conditional cooperativity (12). In contrast, TacAT enhanced *tacAT* promoter activity under the tested conditions, and mutation of OP1 abolished this activation. This regulatory behavior differs from the autorepressive mode described for TacAT-related systems in Salmonella (12), suggesting that homologous TA modules may undergo functional diversification depending on species, genomic context, or regulatory environment. Importantly, our findings do not indicate that TacAT should be classified as a conventional transcription factor. Instead, they demonstrate that a TA-associated complex can acquire promoter regulatory activity, expanding the functional repertoire of TA systems beyond growth inhibition and autorepression.

The requirement for TacT in efficient promoter binding further distinguishes this regulatory system. TacA alone failed to show detectable promoter binding or activation under the tested conditions, whereas the TacA–TacT complex displayed both activities. This observation is consistent with previous studies showing that toxin binding can influence antitoxin conformation, stability, oligomerization, and DNA-binding properties (28). However, the precise mechanistic contribution of TacT remains to be established. TacT may stabilize TacA, facilitate complex formation, or promote a DNA-binding-competent conformation of the TacAT complex. Structural and biochemical analyses, including determination of complex stoichiometry, operator-binding kinetics, and TacA stability, will be necessary to resolve these possibilities.

The downstream targets identified in this study provide a mechanistic explanation for the oxidative stress sensitivity of *tacAT* mutants. ROS-mediated damage includes both oxidative modification and misfolding of proteins, as well as membrane lipid oxidation that compromises envelope integrity and permeability barriers (11). TacAT-induced expression of *clpB* and *htpG* is therefore consistent with a role in restoring proteostasis through protein refolding and disaggregation. Meanwhile, activation of *cadC*, *bhsA*, and *marA* suggests additional functions in maintaining envelope stability and coordinating broader stress adaptation. By targeting these downstream repair processes, TacAT may provide a nonredundant survival advantage during high-level peroxide stress, particularly after the initial ROS burden has been controlled by classical antioxidant systems.

Finally, the attenuation of Δ*tacAT* strains in vivo extends the functional relevance of this regulatory module to infection. In a murine bacteremia model, loss of *tacAT* reduced lethality, bacterial burdens in blood and organs, tissue pathology, and systemic cytokine responses. These phenotypes indicate that TacAT contributes to bacterial persistence and dissemination during systemic infection. Because oxidative stress represents a major host-imposed challenge during infection (24), the peroxide-sensitive phenotype observed in *tacAT* mutants provides a plausible explanation for their reduced fitness in vivo. Nevertheless, systemic infection exposes bacteria to multiple simultaneous stresses, including nutrient limitation, complement-mediated killing, antimicrobial peptides, osmotic stress, envelope stress, and inflammatory immune responses (29). Therefore, the most conservative interpretation is that TacAT functions as a broader stress-adaptive fitness module that enhances *K. pneumoniae* survival under host-associated pressures, with oxidative stress representing a key experimentally supported component.

## Materials and Methods

### Bacterial strains, plasmids, growth conditions and instrument

The *K. pneumoniae* strain used in this study was *CRK3022*, and all mutant strains were generated as described below. *E. coli* DH5α and BL21 strains were obtained from Beijing Qingke Biotechnology Co., Ltd. The plasmids employed included pCDF, pET-22b, pRG970, pBAD, pMDIAI, pFLP-Hyg, and pACBSR-Hyg. Unless otherwise specified, all bacterial strains were cultured in Luria–Bertani (LB) medium. Optical density measurements assays described in this study were performed using a SuPerMax 3500 microplate reader (Shanghai Flash Bio-Tech Co., Ltd.).

### Construction of *K. pneumoniae* mutant strains

*K. pneumoniae* competent cells were prepared on ice. Cultures grown to OD₆₀₀ 0.4–0.6 were harvested at 6,500 rpm for 5 min and washed three times with 10% glycerol. Cells were finally resuspended in glycerol, aliquoted (100 µL), and stored at –80 °C. For transformation, 100 µL of competent cells were mixed with 400 ng DNA or plasmid and electroporated at 2,500 V. LB medium (250 µL) was added post-electroporation, and cells were incubated at 37 °C, 220 rpm for 2 h before plating on selective LB agar.

Targeted gene deletion was performed using pACBSR-Hyg. Wild-type cells were first transformed with pACBSR-Hyg and selected on 100 µg/mL hygromycin B. Competent cells carrying pACBSR-Hyg were then transformed with PCR products containing the upstream and downstream flanking regions of the target gene with an FRT-flanked Apra^R cassette. Transformants were selected on 50 µg/mL apramycin, confirmed by sequencing, and streaked to eliminate the pACBSR-Hyg plasmid.

The Apra^R cassette was excised by introducing pFLP-Hyg into the gene-replaced strain. After selection on hygromycin B and incubation at 43 °C, colonies were screened on LB agar with and without 50 µg/mL ampicillin to isolate marker-free mutants.

### High-Level Oxidative Stress Survival Assay

Overnight cultures of *K. pneumoniae* were diluted 1:100 into fresh medium and grown to mid-exponential phase (OD₆₀₀ ≈ 0.5). To impose a severe oxidative stress challenge, H_o_O_2_ was added to a final concentration of 1%. At the indicated time points (20 min, 1 h, and 2 h) following treatment, aliquots were collected, serially diluted, and plated to determine colony-forming units (CFU). CFU enumeration was used to quantify the fraction of cells that remained viable after prolonged exposure to lethal oxidative stress, thereby assessing bacterial tolerance and survival capacity under extreme oxidative conditions.

### Bioinformatics Analysis

To identify candidate OxyR-responsive loci, the chromosome and plasmid sequences of *K. pneumoniae CRK3022* were obtained from the NCBI database (https://www.ncbi.nlm.nih.gov/). Promoter-proximal noncoding regions upstream of annotated open reading frames were extracted for motif analysis. Unless otherwise specified, promoter-proximal regions were defined as the 100-bp sequences upstream of predicted translational start sites, excluding sequences overlapping adjacent coding regions. Both DNA strands were searched for sequences resembling the *E. coli* OxyR consensus motif ATAG-N6-7-CTAT. Candidate loci were annotated according to the nearest downstream gene or predicted operon. Functional annotations were assigned based on genome annotation and conserved-domain prediction.

To explore potential species-specific differences in OxyR that may underlie divergent regulatory outputs, the amino acid sequences of OxyR from *K. pneumoniae* and *E. coli* were subjected to comparative analyses. Multiple sequence alignment was performed using ESPript (https://espript.ibcp.fr/ESPript/cgi-bin/ESPript.cgi) to visualize conserved residues. Structural models of OxyR from both species were generated using AlphaFold3 (https://alphafoldserver.com/), and structural superposition and comparative analyses were conducted in PyMOL (Schrödinger, LLC) to assess conformational similarities and differences that may contribute to species-specific transcriptional regulation.

### Protein Expression and Purification

*E. coli* BL21 (DE3) strains harboring the pcdf-*tacAT* or pcdf-*oxyR* plasmids were cultured in LB medium at 37 °C with shaking until the optical density at 600 nm (OD₆₀₀) reached approximately 0.8. Protein expression was induced by the addition of IPTG to final concentrations of 0.1 or 0.5 mM, followed by further incubation at 25 °C or 16 °C with shaking overnight.

Cells were harvested by centrifugation and resuspended in lysis buffer containing 15 mM Tris– HCl (pH 8.0), 150 mM NaCl, and 10% (v/v) glycerol. Cell disruption was performed by sonication on ice, and cell debris was removed by centrifugation at 14,000 rpm for 30 min at 4 °C. The clarified supernatant was applied to a Ni²⁺-affinity column, and bound proteins were eluted with buffer containing 300 mM imidazole. The eluted proteins were further purified using an ÄKTA purification system (GE Healthcare).

### EMSA

The promoter regions of the target genes were amplified from the genomic DNA of *K. pneumoniae* strain *CRK3022*. For DNA–protein binding assays, purified proteins were incubated with the corresponding promoter DNA fragments at varying molar ratios in a total reaction volume of 20 μL. Binding reactions were performed in buffer containing 15 mM Tris–HCl (pH 8.0), 150 mM NaCl, and 10% (v/v) glycerol.

The reaction mixtures were incubated at 4 °C for 30 min and subsequently resolved by electrophoresis on 8% native polyacrylamide gels in 1× TBE (Tris–borate–EDTA) buffer at 120 V for 1–3 h. After electrophoresis, the gels were stained with ethidium bromide (EB), and DNA– protein complexes were visualized using a Bio-Rad gel imaging system.

### RNA extraction and real-time quantitative PCR (RT-qPCR)

Total RNA was extracted using TRIzol reagent (Vazyme, China) according to the manufacturer’s instructions. Briefly, bacterial cells were lysed in TRIzol, followed by chloroform extraction and isopropanol precipitation. The RNA pellet was washed with 75% ethanol and dissolved in RNase-free water. RNA concentration and purity were measured using a NanoDrop spectrophotometer (Thermo Scientific).

Complementary DNA (cDNA) was synthesized from total RNA using a HiScript reverse transcription kit (Vazyme, China) following the manufacturer’s protocol. RT-qPCR was performed using ChamQ SYBR qPCR Master Mix (Vazyme, China) on a real-time PCR system. Relative gene expression levels were normalized to rpoD and calculated using the 2⁻ΔΔCt method. All experiments were performed with at least three biological replicates.

### β-galactosidase assay

The promoter regions (∼200 bp upstream of the start codon) of the genes of interest were cloned upstream of the promoterless lacZ gene in the pRG970-KM vector to generate lacZ transcriptional reporter constructs. The resulting reporter plasmids, together with the corresponding protein expression plasmids, were co-transformed into *E. coli*. Transformants were grown in LB medium at 37 °C to an OD₆₀₀ of approximately 0.6.

For the β-galactosidase assay, 1 mL of culture was harvested and permeabilized with 50 μL of 0.1% sodium dodecyl sulfate (SDS). The reaction was initiated by adding 40 μL of o-nitrophenyl-β-D-galactopyranoside (ONPG) solution (4 mg/mL). The reaction was allowed to proceed until a visible yellow color developed, and the reaction time was recorded for Miller unit calculation; reaction times ranged from 5 s to 6 min depending on signal intensity. β-Galactosidase activity was determined by measuring the absorbance at 420 nm and 550 nm, reflecting the enzymatic conversion of the colorless ONPG substrate into the yellow product o-nitrophenol. β-Galactosidase activity was calculated in Miller units using the formula:

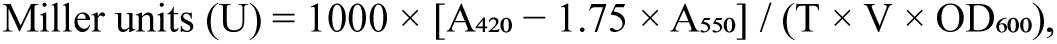

where T represents the reaction time (min) and V denotes the volume of culture used in the assay (mL).

### Total Protein Extraction

Frozen samples were kept on ice and resuspended in lysis buffer containing 8 M urea, 1% SDS, and protease inhibitors. Samples were homogenized using a high-flux tissue grinder for three cycles of 180 s each, followed by non-contact cryogenic sonication for 30 min. Lysates were centrifuged at 16,000 × g at 8 °C for 30 min, and protein concentrations in the supernatants were determined using the BCA Protein Assay Kit (Thermo Scientific) according to the manufacturer’s instructions. Extracted proteins were subsequently analyzed by SDS-PAGE.

### Protein digestion

For enzymatic digestion, 100 μg of protein was resuspended in triethylammonium bicarbonate (TEAB) buffer to a final concentration of 100 mM. Proteins were reduced with tris(2-carboxyethyl)phosphine (TCEP) at a final concentration of 10 mM at 37 °C for 60 min, followed by alkylation with iodoacetamide (IAM) at a final concentration of 40 mM for 40 min at room temperature in the dark. After centrifugation at 10,000 × g at 4 °C for 20 min, the pellet was collected and resuspended in 100 μL of 100 mM TEAB buffer. Trypsin was then added at a trypsin-to-protein mass ratio of 1:50, and digestion was carried out at 37 °C overnight.

### Peptide Desalting and Quantification

Following tryptic digestion, peptides were dried under vacuum. The dried peptides were subsequently reconstituted in 0.1% trifluoroacetic acid (TFA) and desalted using HLB solid-phase extraction cartridges. After desalting, peptides were dried again in a vacuum concentrator. Peptide concentrations were then determined by measuring UV absorbance using a Nano Drop One spectrophotometer (Thermo Scientific).

### DIA mass detection

Based on peptide quantification results, peptides were analyzed using a Vanquish Neo UHPLC system coupled to an Orbitrap Astral mass spectrometer (Thermo Fisher Scientific, USA) at Majorbio Bio-Pharm Technology Co., Ltd. (Shanghai, China). Peptide separation was performed on a uPAC High Throughput column (75 μm × 5.5 cm; Thermo Fisher Scientific) using solvent A (water containing 2% acetonitrile and 0.1% formic acid) and solvent B (water containing 80% acetonitrile and 0.1% formic acid). The total chromatographic gradient was set to 8 min. Data-independent acquisition (DIA) was carried out on the Orbitrap Astral mass spectrometer operated in DIA mode. The MS scanning range was set from m/z 100 to 1700.

### Protein identification

DIA raw data were processed and searched using Spectronaut software (version 19). The search parameters were set as follows: peptide length was restricted to 7–52 amino acids; trypsin/P was specified as the proteolytic enzyme with a maximum of two missed cleavage sites allowed. Carbamidomethylation of cysteine residues was defined as a fixed modification, whereas oxidation of methionine and protein N-terminal acetylation were set as variable modifications.

Protein and peptide identifications were filtered using a false discovery rate (FDR) threshold of ≤1% at both the protein and peptide levels. Only peptides with a confidence score ≥99% were retained for further analysis. The extracted ion chromatogram (XIC) width was set to ≤75 ppm. Protein quantification was performed using the MaxLFQ algorithm.

### Statistical analyses

Bioinformatic analysis of the proteomic data was performed using the Majorbio Cloud platform (https://cloud.majorbio.com). Statistical analyses were conducted in R to calculate fold changes (FC) and raw P values for pairwise comparisons between groups using a two-sided t-test. The Benjamini–Hochberg method was applied for multiple-testing correction to control the false discovery rate (FDR). Differential proteins were defined as those with FC > 1.5 and adjusted P < 0.05.

Functional annotation of all identified proteins was carried out based on Gene Ontology (GO) terms (http://geneontology.org/). GO enrichment analysis was subsequently performed using the identified DEPs to evaluate the overrepresentation of functional categories, with particular emphasis on biological process annotations.

### Mouse Bacteremia Model

Overnight bacterial cultures were diluted 1:100 into fresh medium and grown to mid-exponential phase (OD₆₀₀ ≈ 0.5). Cells were harvested by centrifugation, washed three times with phosphate-buffered saline (PBS), and resuspended in PBS. Bacterial suspensions were adjusted to the desired concentrations and administered to mice via tail vein injection at either a high dose (5 × 10⁶ CFU) or a low dose (1 × 10⁵ CFU) to establish the bacteremia model.

### Detection of Bacterial Survival in Mice

Following establishment of the bacteremia model using a low bacterial dose, bacterial survival in mice was assessed at 24 h and 48 h post-infection. Blood samples were collected via tail vein and serially diluted in phosphate-buffered saline (PBS) before plating on LB agar plates. After incubation at 37 °C for 16 h, colony-forming units (CFU) were enumerated.

At 48 h post-infection, mice were euthanized and major organs, including the lungs, liver, and spleen, were aseptically harvested. Tissues were homogenized in PBS, and the homogenates were serially diluted and plated on LB agar plates. After incubation at 37 °C for 16 h, bacterial loads in each organ were determined by CFU counting.

### Hematoxylin and Eosin (H&E) Staining

To evaluate tissue inflammation and pathological damage, mice subjected to low-dose infection were euthanized at 48 h post-infection, and major organs, including the lungs and liver, were aseptically collected. Tissue samples were gently rinsed with phosphate-buffered saline (PBS) to remove residual blood and debris and subsequently fixed in 4% paraformaldehyde at room temperature for 24 h.

After fixation, tissues were dehydrated through a graded ethanol series, embedded in paraffin, and sectioned into 4–5 μm-thick slices. Tissue sections were then subjected to standard hematoxylin and eosin (H&E) staining procedures. Briefly, sections were deparaffinized, rehydrated, stained with hematoxylin to visualize nuclei, and counterstained with eosin to highlight cytoplasmic and extracellular structures.

Stained sections were examined using a light microscope, and representative images were captured to assess inflammatory cell infiltration, tissue architecture disruption, and pathological alterations associated with bacterial infection.

### Cytokine Quantification

To quantify inflammatory cytokine levels, including IL-6, TNF-α, IL-10, and CXCL1, peripheral blood samples were collected from mice at 48 h after low-dose infection. Blood samples were allowed to clot at room temperature and were subsequently centrifuged at 3,000 × g for 10 min to obtain serum. Serum samples were stored at −80 °C until analysis.

Cytokine concentrations were measured using commercially available enzyme-linked immunosorbent assay (ELISA) kits (Lianke Biotechnology, China) according to the manufacturer’s instructions. Briefly, standards and serum samples were added to antibody-coated 96-well plates and incubated at 37 °C. After washing to remove unbound components, biotinylated detection antibodies and horseradish peroxidase (HRP)-conjugated streptavidin were sequentially added. Following substrate incubation, the reaction was terminated with stop solution, and absorbance was measured at 450 nm using a microplate reader.Cytokine concentrations were calculated based on standard curves and expressed as picograms per milliliter (pg/mL).

### Statistics and reproducibility

All data were analyzed using GraphPad Prism software and are presented as mean ± standard error of the mean (SEM). All experiments were performed with at least three independent biological replicates. Two-tailed Student’s t-tests were used for comparisons of gene expression levels, β-galactosidase reporter activity, bacterial burdens (after log₁₀ transformation), and cytokine measurements. Survival curves from mouse infection experiments were analyzed using the Mantel– Cox (log-rank) test. Statistical significance was defined as P < 0.05.

### Data availability

Proteomics raw data have been deposited in the China National Center for Bioinformation under accession number OMIX014779.

## Acknowledgements

This work was supported by the National Natural Science Foundation of China (32370187 to Rui Bao) and the China Postdoctoral Science Foundation (GZC20241164 and 2025M771483 to Ninglin Zhao; GZC20241162 and 2025HXBH108 to Xingyu Mou). We also thank the CAST Youth Science and Technology Talent Cultivation Program for Doctoral Students for support.

## Author contributions

J.Z., H.C.M. and Z.S. conceived and designed the experiments; J.Z., H.C.M., Z.S., S.Z.L., J.Y.R., N.L.Z. and X.Y.M. performed the experiments; J.Z., H.C.M., Z.S. and C.Y.C. processed data analysis; J.Z., H.C.M., Z.S., H.L., H.X.L., H.B.W. and R.B. wrote the manuscript and supervised the overall study.

## Supplementary information

**Figure S1.**
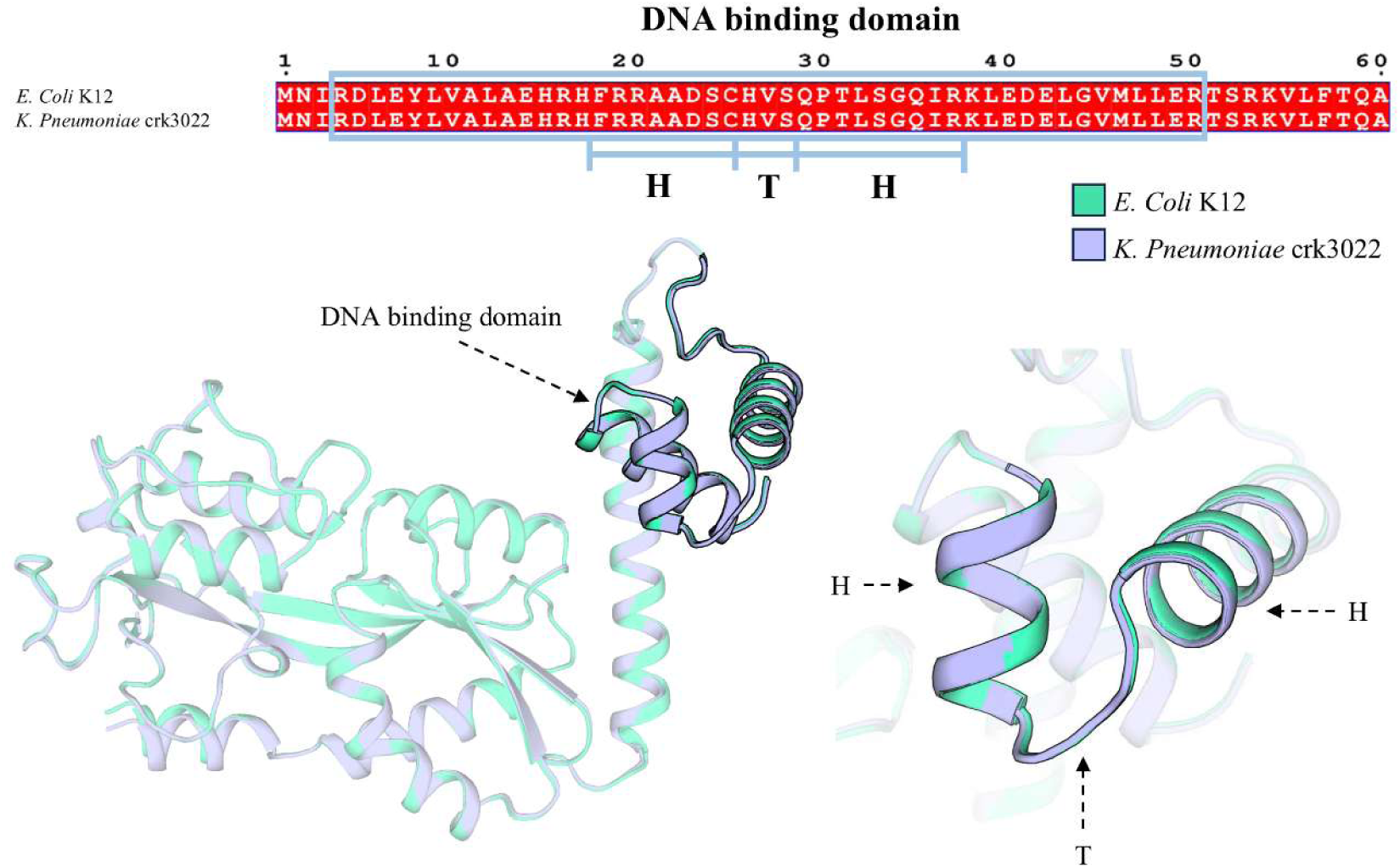
Conservation of the OxyR DNA-binding region between *K. pneumoniae CRK3022* and *Escherichia coli* K-12. The helix-turn-helix (HTH) region is highly conserved, supporting the use of the *E. coli* OxyR consensus motif for candidate motif screening. Structural superposition of predicted OxyR models revealed high overall similarity, with a global RMSD of 0.265 Å and a DNA-binding domain RMSD of 0.049 Å. The *E. coli* K-12 OxyR model is shown in light green, whereas the *CRK3022* OxyR model is shown in purple.

**Figure S2.**
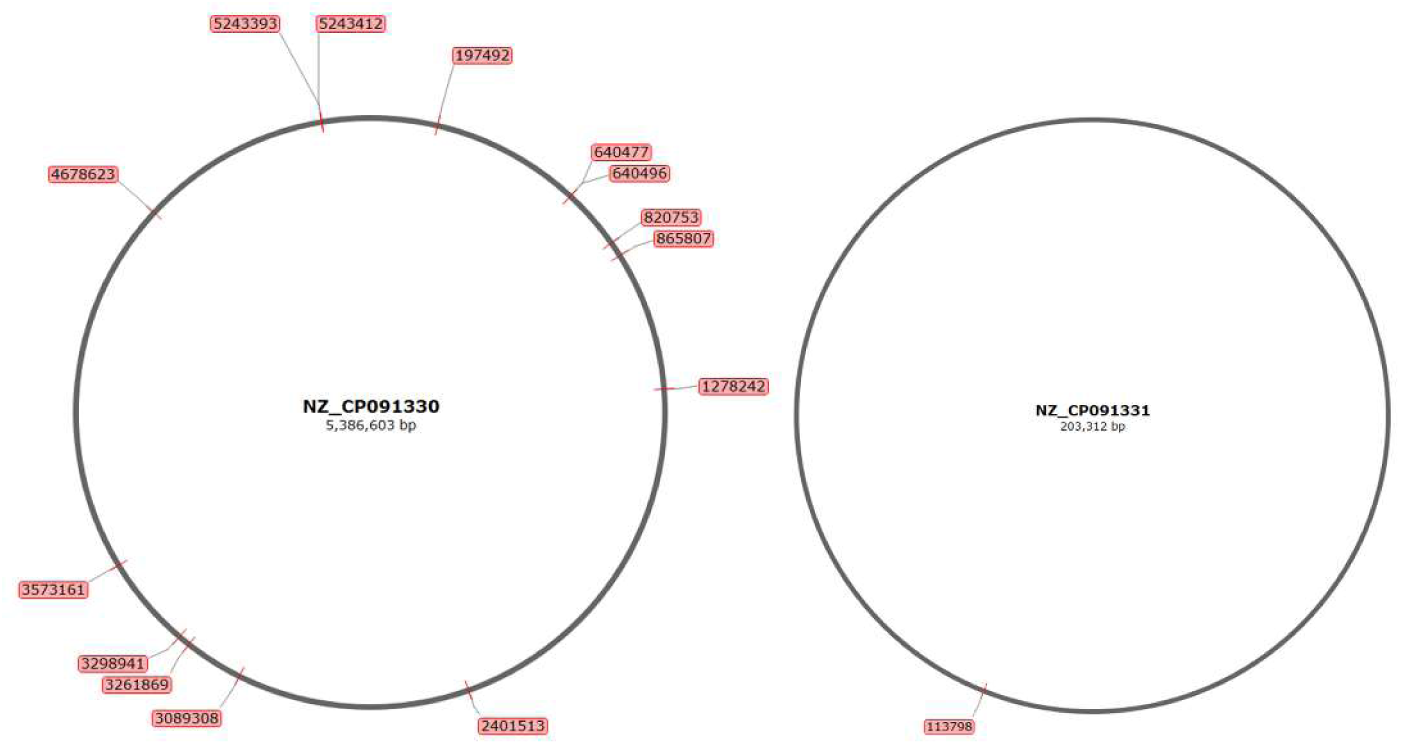
Distribution of predicted OxyR-binding sites in Klebsiella pneumoniae *CRK3022*. NZ_CP091330 represents the chromosome, where 14 predicted OxyR-like sites were identified, whereas NZ_CP091331 represents plasmid 1, containing one additional predicted OxyR-like site.

**Figure S3.**
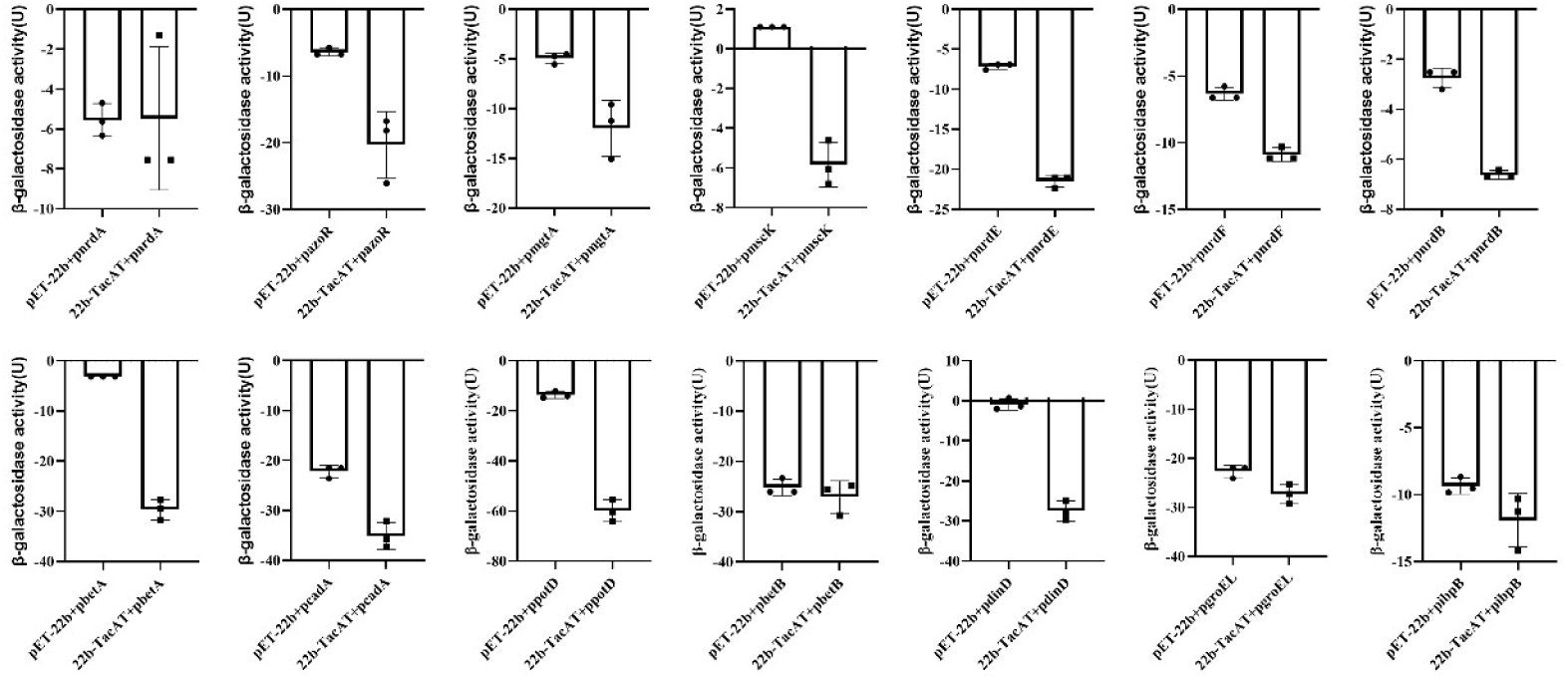
β-Galactosidase reporter assays of additional stress-related genes upregulated in the proteomic comparison between Δ*tacAT*-pBAD-*tacAT* and Δ*tacAT*-pBAD strains. These genes did not show significant TacAT-dependent promoter activation under the tested conditions and were therefore not classified as candidate direct TacAT transcriptional targets.

**Table S1.**
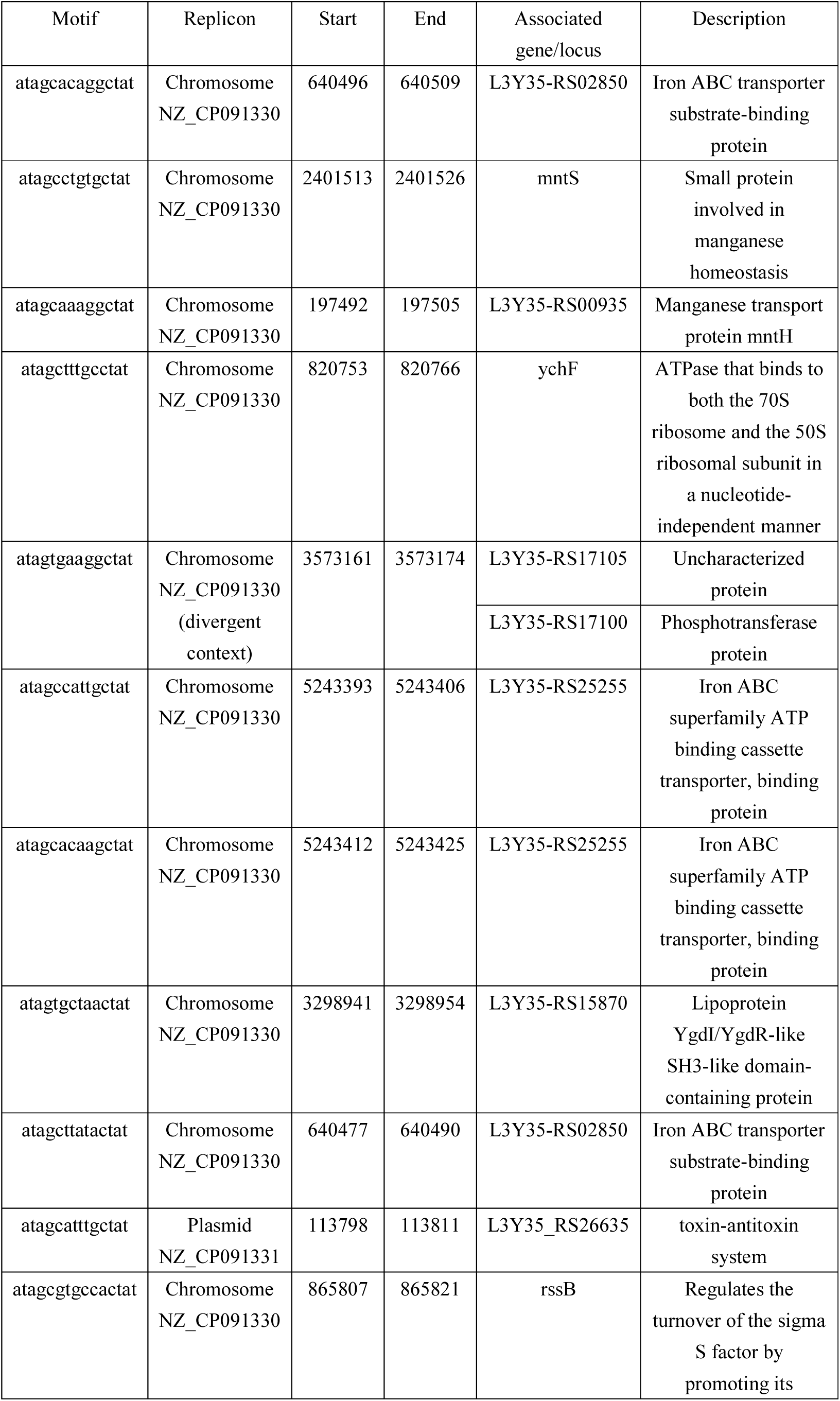
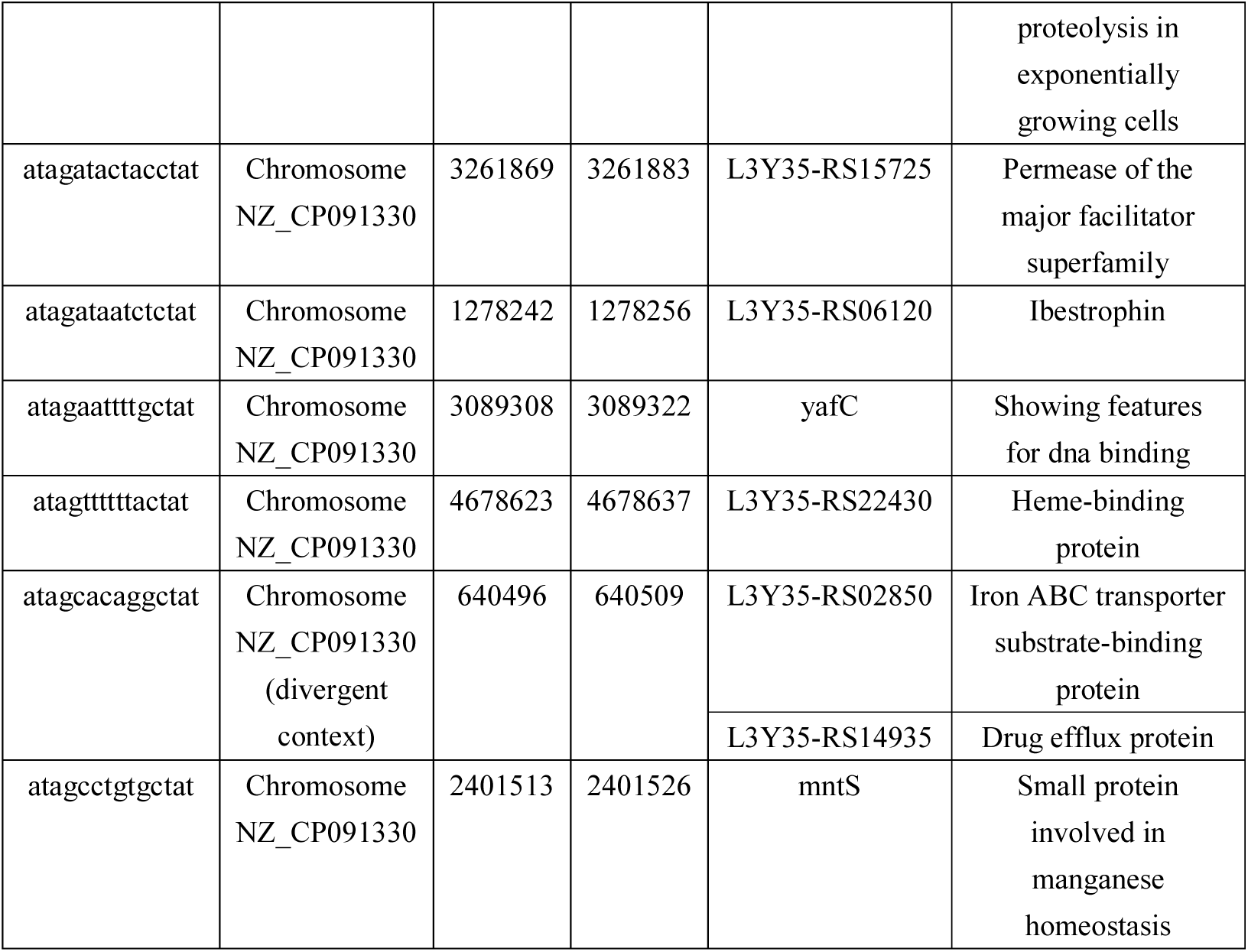
Predicted OxyR-binding sites and associated genes in *K. pneumoniae CRK3022*.

**Table S2.** Stress-related genes among upregulated proteins identified by proteomic analysis comparing Δ*tacAT*–pBAD-*tacAT* and Δ*tacAT*–pBAD strains.

| Accession | Symbol | Protein Name | Log2FC | P_value | Functional category |
| --- | --- | --- | --- | --- | --- |
| A0A086IXC1 | nrdA | Ribonucleoside-diphosphate reductase | 0.7807 | 0.000209 | DNA synthesis / stress response |
| A0A0C7KDY0 | azoR | FMN dependent NADH: quinone oxidoreductase | 0.6713 | 7.55E-05 | Redox homeostasis |
| A0A086ICS8 | mgtA | Magnesium-transporting ATPase, P-type 1 | 0.8935 | 0.008847 | Metal ion homeostasis |
| A0A1Y0Q1F3 | mscK | Mechanosensitive channel MscK | 0.774 | 0.04776 | Envelope/membrane adaptation |
| A0A378BWB1 | nrdE | Ribonucleoside-diphosphate reductase | 0.821 | 6.40E-05 | DNA synthesis / stress response |
| A0A377TXU5 | nrdF | ribonucleoside-diphosphate reductase | 0.9247 | 0.000645 | DNA synthesis / stress response |
| A0A377ZB73 | nrdB | ribonucleoside-diphosphate reductase | 0.9159 | 0.00204 | DNA synthesis / stress response |
| A0A0H3GL29 | betA | Oxygen-dependent choline dehydrogenase | 1.0155 | 0.000174 | Osmotic stress adaptation |
| A0A377Z5U7 | cadC | DNA-binding transcriptional activator CadC | 0.6537 | 0.004979 | Stress-responsive regulation |
| A0A086IBI2 | cadA | Inducible lysine decarboxylase | 0.9197 | 0.000349 | Acid/envelope stress adaptation |
| A0A4S7WIB9 | potD | Putrescine-binding periplasmic protein | 0.7685 | 0.03055 | Transport / stress adaptation |
| A0A060VM76 | betB | Betaine aldehyde dehydrogenase | 1.0293 | 6.08E-05 | Osmotic stress adaptation |
| A0A0C7KE20 | dinD | DNA damage-inducible protein D | 1.2339 | 1.46E-05 | DNA damage response |
| H6VX41 | groEL | GroEL | 1.6743 | 0.01399 | Protein quality control |
| A0A378AAM7 | clpB | Protein disaggregation chaperone | 11.5406 | 2.02E-05 | Protein quality control |
| A0A081ISF9 | bhsA | DUF1471 domain-containing protein | 1.0551 | 0.001648 | Envelope stress |
| A0A080SXB9 | marA | Multiple antibiotic resistance protein MarA | 0.9333 | 0.02798 | Envelope/membrane adaptation |
| A0A483LKY1 | htpG | Chaperone protein HtpG | 1.1268 | 0.0228 | Protein quality control |
| A0A0C7KDK9 | ibpB | Small heat shock<br>protein IbpB | 1.1791 | 5.96E-06 | Protein quality<br>control |

**Table S3.**
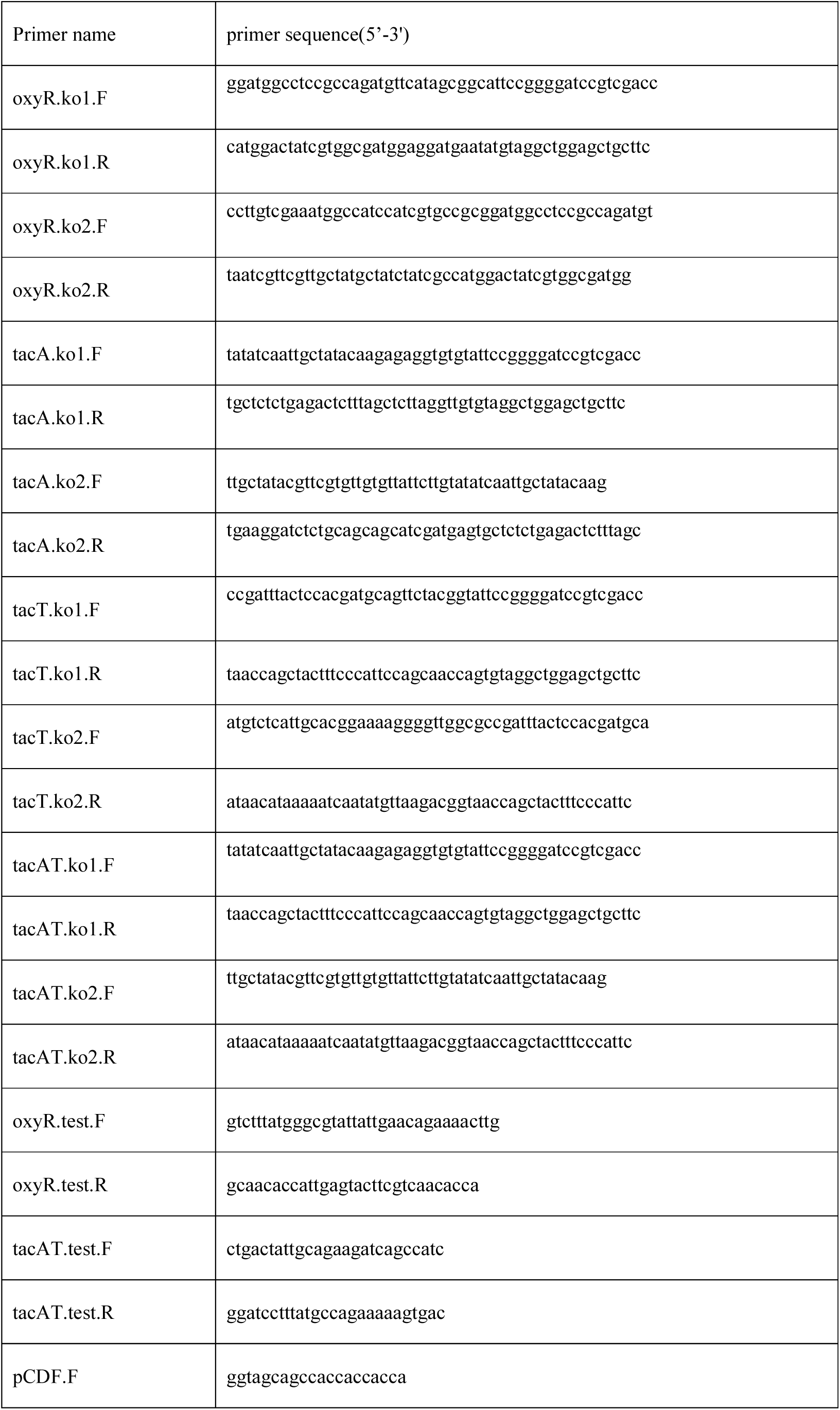
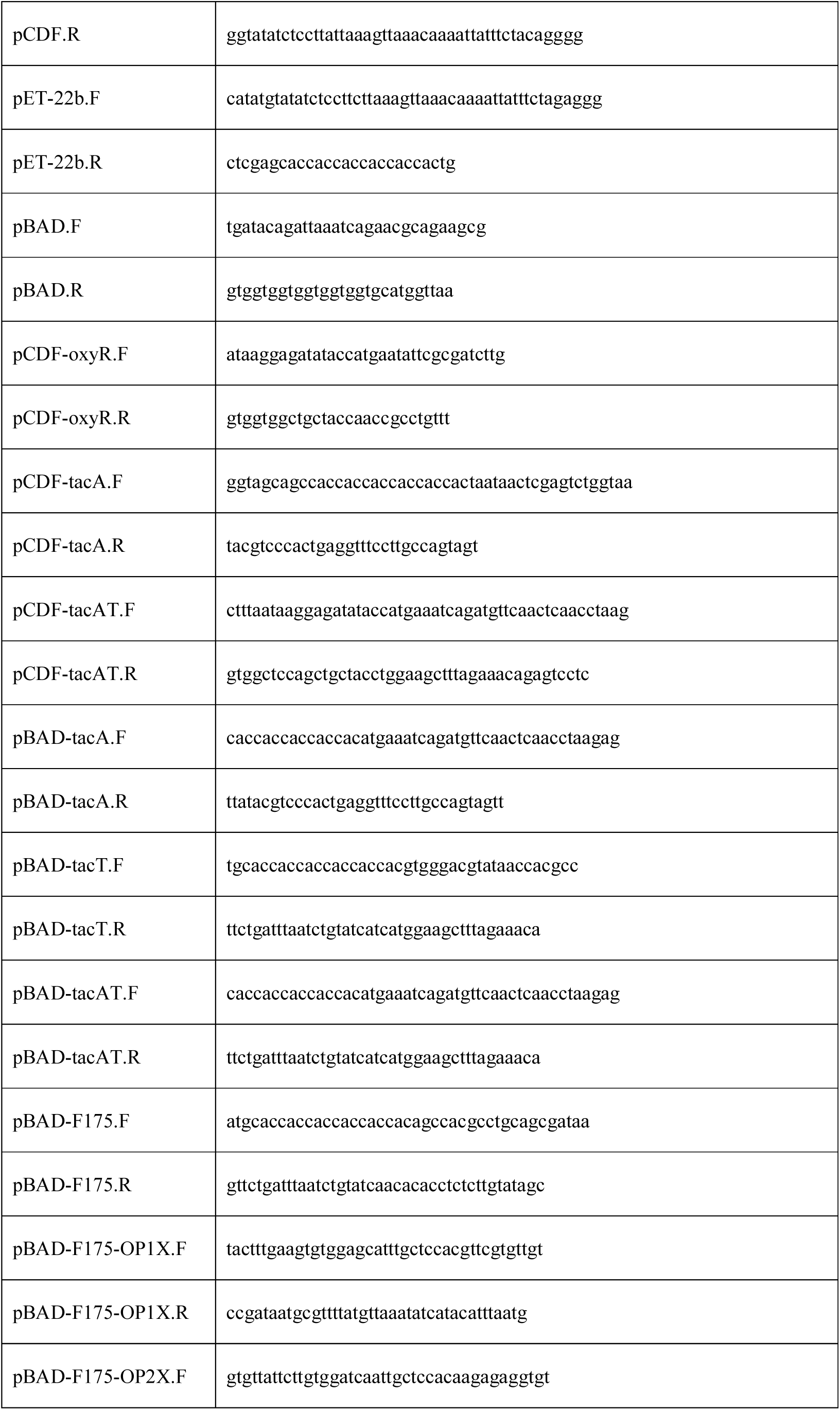
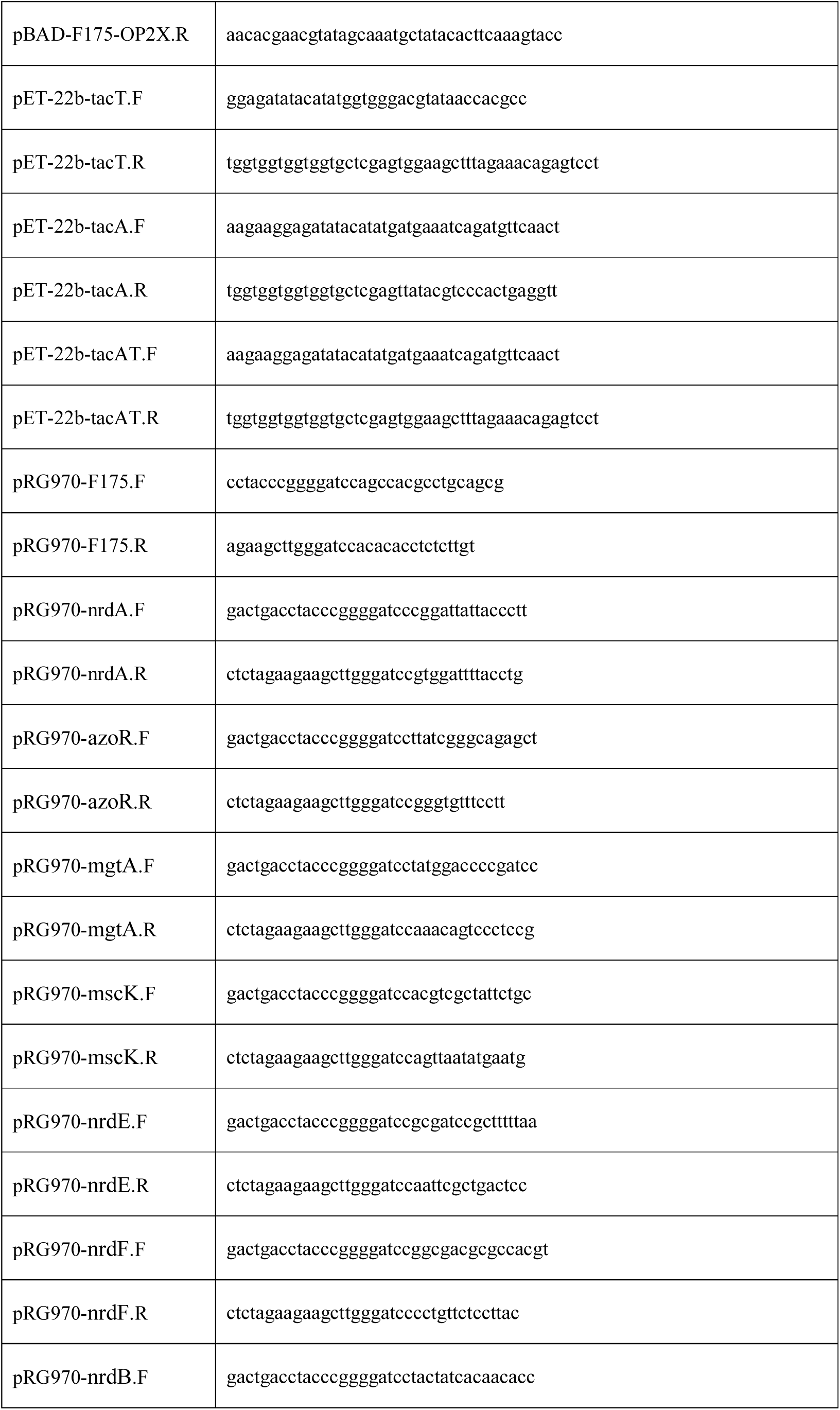
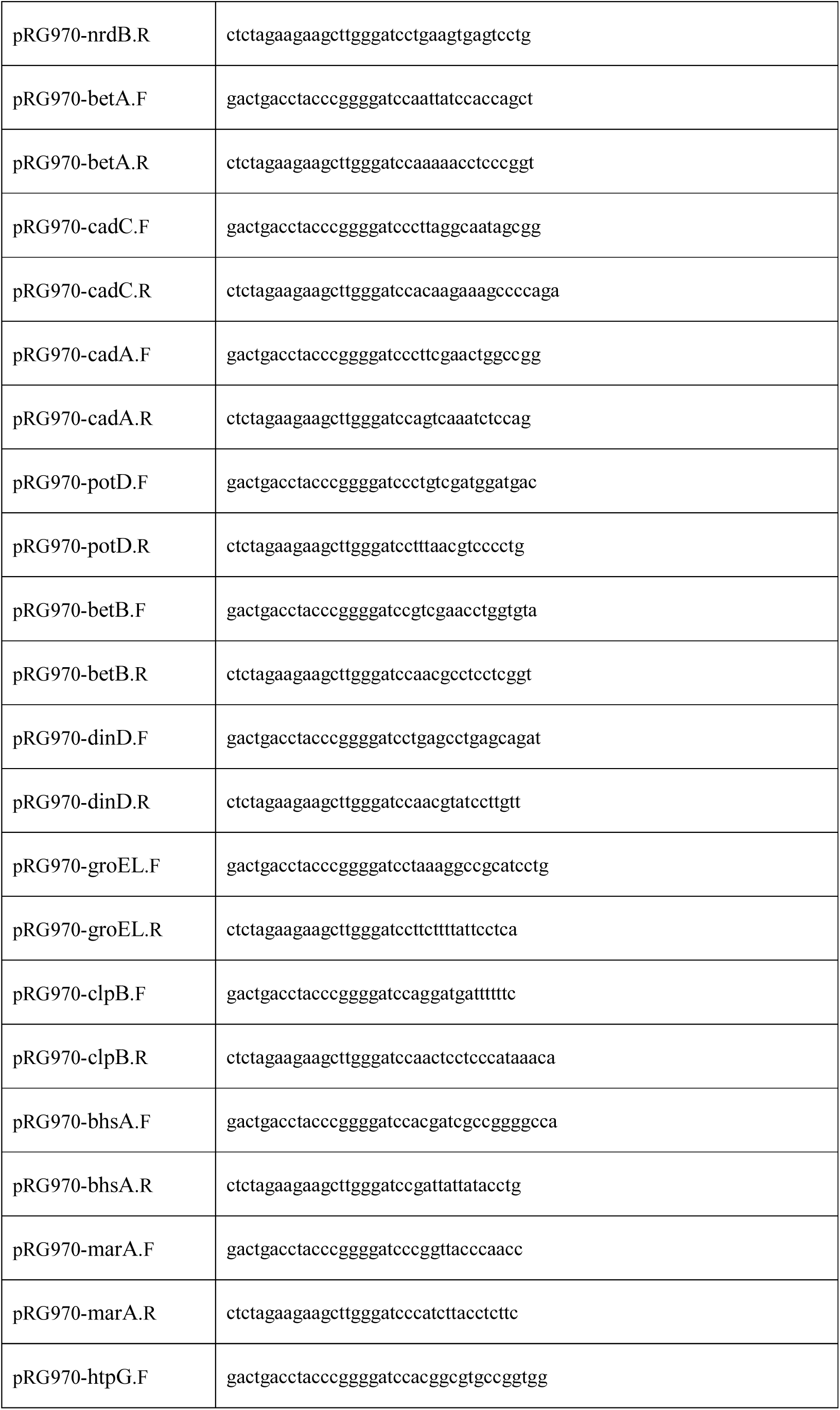
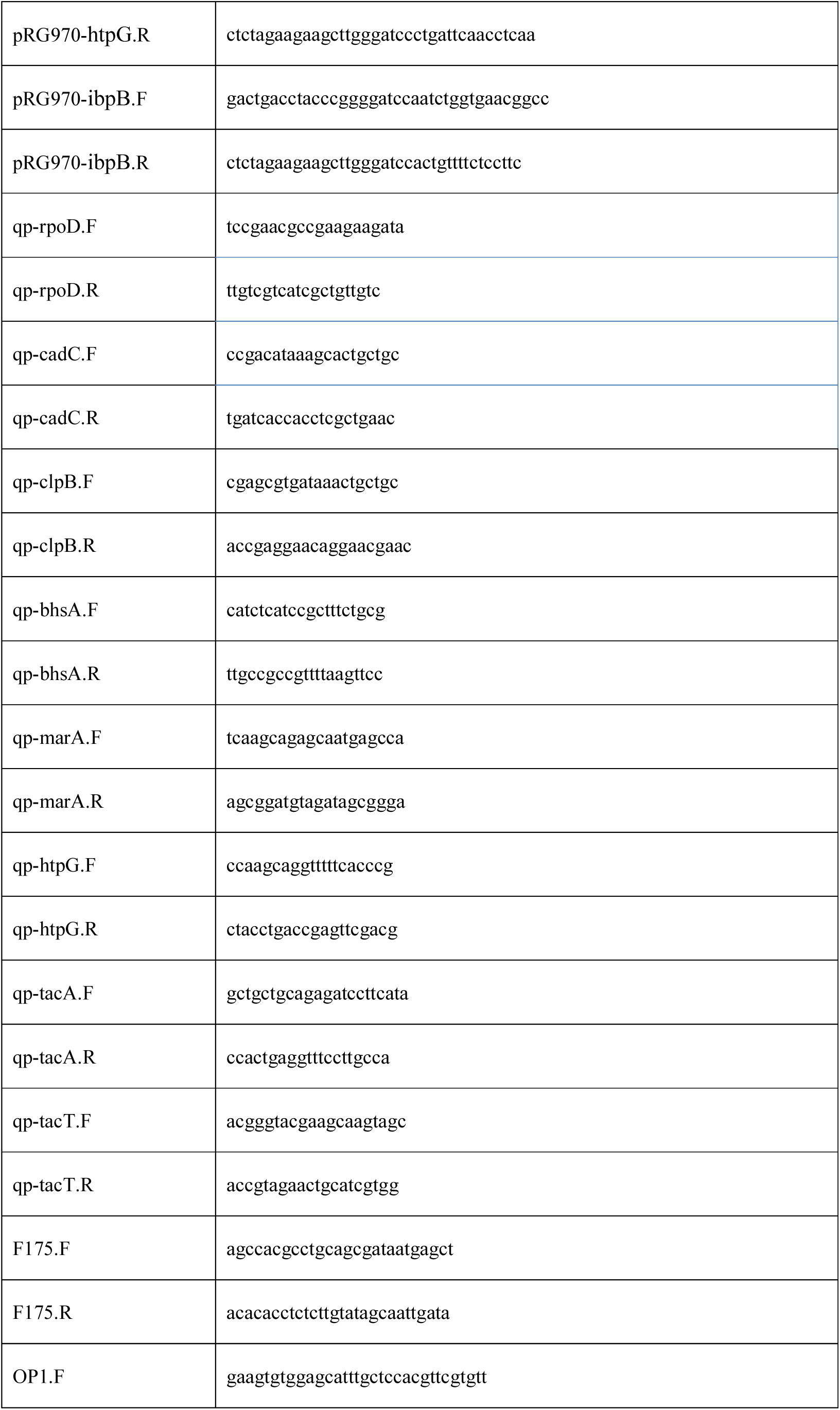
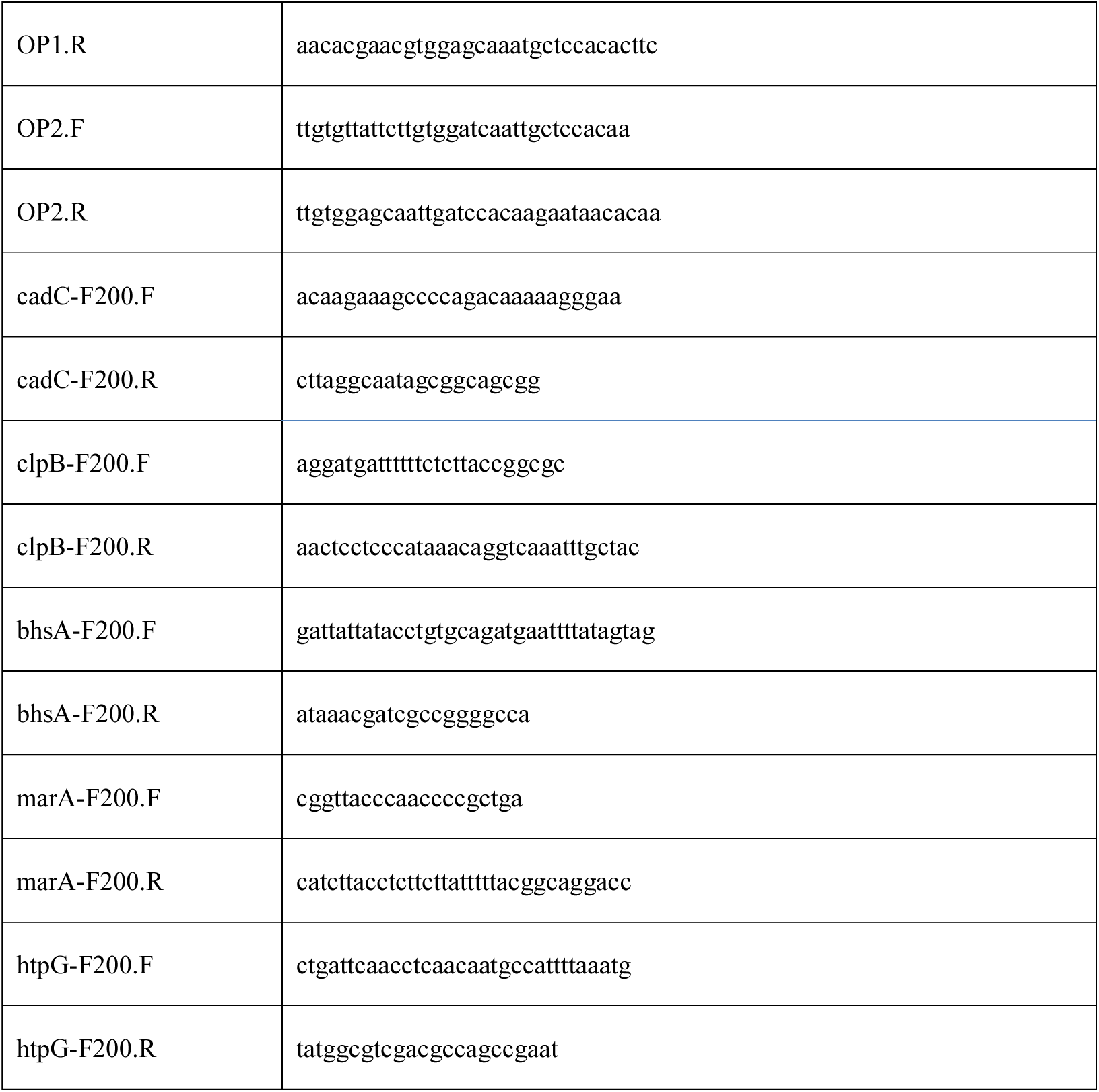
All primers used in this study.

